# Bat RNA viruses employ viral RHIMs orchestrating species-specific cell death programs linked to Z-RNA sensing and ZBP1-RIPK3 signaling

**DOI:** 10.1101/2023.08.23.554434

**Authors:** Sanchita Mishra, Disha Jain, Ayushi Amin Dey, Sahana Nagaraja, Mansi Srivastava, Oyahida Khatun, Keerthana Balamurugan, Micky Anand, Shashank Tripathi, Mahipal Ganji, Sannula Kesavardhana

**Affiliations:** Department of Biochemistry, Division of Biological Sciences, Indian Institute of Science, Bengaluru, Karnataka 560012, India; Department of Microbiology and Cell Biology, Division of Biological Sciences, Indian Institute of Science, Bengaluru 560012, India; Centre for Infectious Disease Research, Indian Institute of Science, Bengaluru 560012, India

**Author notes:** <u>Correspondence:</u> Sannula Kesavardhana Department of Biochemistry Division of Biological Sciences, Indian Institute of Science (IISc), Bengaluru, Karnataka 560012, India. Equal contribution (Disha Jain, Ayushi Amin Dey, Sahana Nagaraja).

**Keywords:** Bat viruses, viral RHIMs, SARS-CoV-2, coronavirus, Nsp13, cell death, bats, Z-RNA, ZBP1, necroptosis, RIPK3, zoonotic viruses, virus adaptation, inflammation, host defense, innate immunity

## Abstract

RHIM is a protein motif in cell death proteins that assembles higher-order signaling complexes and triggers regulated cell death, which in itself limits virus spread and additionally triggers inflammation for mounting immune responses. A few DNA viruses employ viral RHIMs mimicking host RHIMs. However, these viral RHIMs counteract host cell death by interacting with host RHIM proteins and blocking complex formation to alleviate antiviral defenses. Whether RNA viruses operate such viral RHIMs remains unknown. RHIM-protein signaling promotes lung damage and cytokine storm in respiratory RNA virus infections, arguing the presence of viral RHIMs. Here, we report the novel viral RHIMs in Nsp13 and Nsp14 of SARS-CoV-2 and other bat RNA viruses, providing the basis for bats as the hosts for their evolution. Nsp13 promoted cell death in bat and human cells, however, viral RHIM of Nsp13 is more critical for human cell death than bat cells, suggesting species-specific regulation. The conformation of RNA-binding channel in Nsp13 is critical for cell death in bat and human cells. Nsp13 showed RHIM-dependent interactions with ZBP1 and RIPK3 and promoted the formation of large insoluble complexes of ZBP1 and RIPK3. Also, Nsp13 promoted ZBP1-RIPK3 signaling-mediated cell death dependent on intracellular RNA ligands. Intriguingly, the SARS-CoV-2 genome consists of bona fide Z-RNA-forming segments. These SARS-CoV-2 Z-RNA segments promoted Nsp13-dependent cell death, further revealing Nsp13’s association with Z-RNA sensing and ZBP1-RIPK3 signaling. Our findings reveal the functional viral RHIMs of bat-originated RNA viruses regulating host cell death associated with Z-RNA sensing and ZBP1-RIPK3 signaling activation. These observations allow the understanding of mechanisms of cellular damage and cytokine storm in SARS-CoV-2 and other bat-originated RNA virus infections.

**GRAPHICAL ABSTRACT:** 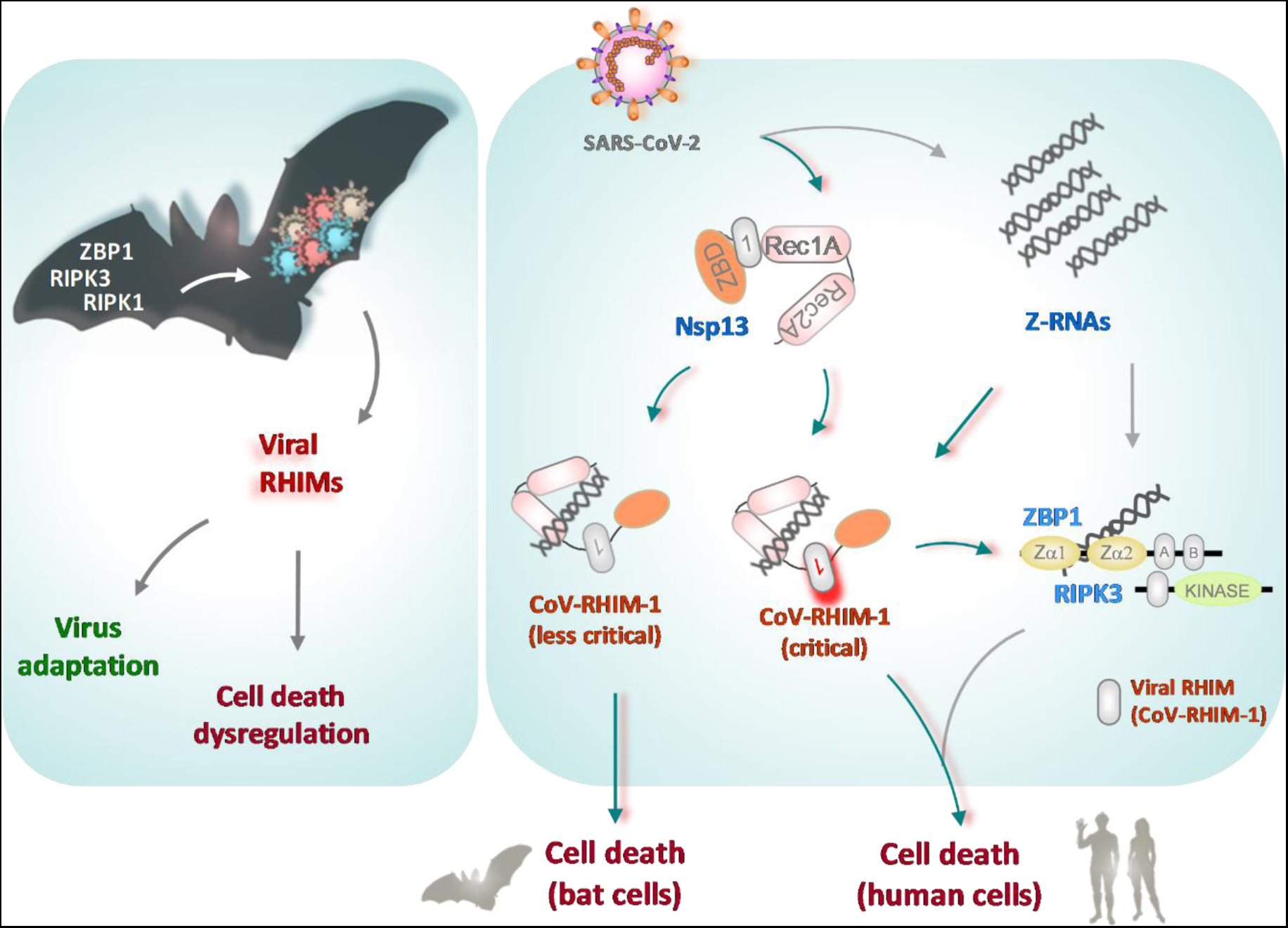

**One-sentence summary:** Bat-associated RNA viruses employ viral RHIMs and regulate host cell death.

## INTRODUCTION

Pathogenic RNA virus infections often result in uncontrolled tissue damage and inflammatory responses, which prime disease pathogenesis (*1–3*). Severe acute respiratory syndrome coronavirus 2 (SARS-CoV-2) is associated with mild to severe respiratory infections and causes coronavirus disease 2019 (COVID-19) (*4, 5*). COVID-19 patients develop pneumonia and acute respiratory distress syndrome (ARDS), which lead to the failure of respiratory function (*1, 4, 5*). However, the host and viral mechanisms underlying disease pathogenesis in COVID-19 and other human respiratory infections are unclear. Aberrant cell death and cytokine storm have been reported to be associated with COVID-19 pathogenesis (*1, 2, 6*). RNA viruses, including CoVs, are adapted to evade host defense responses to promote their spread and pathogenesis in the infected hosts (*7, 8*). Also, these RNA viruses show restricted type I interferon (IFN) production and cause dysregulated immune and proinflammatory responses in the lung (*1, 8, 9*).

Regulated inflammatory cell death is an innate host defense response to virus infections (*3, 10, 11*). Inflammatory cell death removes the viral replication niche by eliminating the virus-infected cells and mounts protective immune responses and repair processes (*1, 9*). Type I IFNs promote inflammatory cell death as a protective host defense mechanism during viral infections (*9, 12*). Human receptor-interacting serine/threonine protein kinase 1 (RIPK1), RIPK3, Toll/IL-1 receptor domain-containing adapter protein-inducing IFN-β (TRIF), and Z-nucleic acid binding protein 1 (ZBP1) are inflammatory cell death signaling proteins (*13, 14*). These proteins have a conserved receptor-interacting protein (RIP)-homotypic interaction motif (RHIM), which confers protein-protein interactions of these cell death proteins. RHIM-mediated protein-protein interactions trigger the formation of higher-order signaling complexes, structurally resembling an amyloid fibrillar assembly, which promote cell death and inflammation (*14–16*). The RIPK3 in this complex mediates the activation of mixed lineage kinase domain-like (MLKL), leading to the execution of an inflammatory form of cell death necroptosis (*15, 17, 18*). The RIPK3 also promotes RHIM-dependent recruitment of RIPK1 and activation of caspase-8-dependent apoptosis (*10, 18, 19*). Necroptosis is essential for eliminating virus-infected cells and blocking viral spread, but its dysregulation sometimes aggravates inflammation and tissue damage (*3, 9–11*). RIPK3 and ZBP1 are the key triggers of necroptosis during viral infections (*19–27*). These proteins are also associated with activating inflammasome-driven pyroptosis, and a multifaceted cell death modality called PANoptosis (*6, 10, 28, 29*). ZBP1 is an innate immune sensor recognizing viral Z-RNAs generated during viral replication (*3, 10, 30*). Upon ZBP1 activation, RHIM-mediated interaction of ZBP1 and RIPK3 triggers cell death and inflammation. Due to the critical role of RHIMs in driving ZBP1-RIPK3 signaling and antiviral host defense, a few DNA viruses (herpesviruses) have evolved to mimic host RHIMs to counteract ZBP1-RIPK3 signaling (*20, 24, 31*). These viral RHIMs bind to host RIPK3 to restrict its interaction with RIPK1 and ZBP1 and the formation of cell death signaling complexes to help continue viral replication and spread (*20, 21, 24, 26, 31–33*). Interestingly, viral RHIMs of HSV-1 restrict RIPK3-mediated cell death in human cells, which is a natural host species, but promote cell death in mouse cells, suggesting the species-specific functions of viral RHIMs in either promoting or restricting cell death activation (*24, 31, 34*). Recent studies show that poxvirus vaccinia encodes Z-RNA competitors of ZBP1 activation to regulate ZBP1-RIPK3 signaling (*35, 36*). Unlike DNA viruses, influenza viruses do not encode any RHIM-mediated suppressors but instead activate ZBP1-RIPK3-dependent cell death with inflammatory damage (*1, 3, 10*). Also, aberrant cell death, inflammation, and necrotic lung damage are the clinical signs observed during pathogenic influenza and SARS-CoV-2 infections (*1, 6, 28, 37, 38*). This indicates a possible lack of viral RHIMs or the presence of cell death-promoting viral RHIMs in pathogenic respiratory viruses. However, whether RNA viruses operate viral RHIMs to modulate host cell death responses is unknown.

Here, we report the identification of novel viral RHIMs of CoVs (CoV-RHIMs) in Nsp13 and Nsp14 proteins. CoV-RHIM in Nsp13 of pathogenic CoVs (SARS-CoV, MERS-CoV, and SARS-CoV-2) is highly conserved and retains RHIM-like features compared to other human-infected CoVs, whereas CoV-RHIMs in Nsp14 were conserved in less pathogenic human CoVs. Also, our observations indicate bat RHIM-proteins in driving the evolution of viral RHIMs in RNA viruses. Unlike RHIM proteins of DNA viruses, CoV-RHIM of SARS-CoV-2 Nsp13 promoted cell death and the RNA binding channel of Nsp13 is critical for this function. Intriguingly, Nsp13 showed species-specific distinct mechanisms of cell death regulation. Also, SARS-CoV-2 Nsp13 formed RHIM-dependent complexes with the ZBP1 and promoted the formation of ZBP1 and RIPK3 higher-order complexes. Further biochemical studies suggest the role of SARS-CoV-2 Z-RNAs in promoting Nsp13-mediated cell death. Our observations demonstrate the evolution and operation of viral RHIMs in RNA viruses that might be linked to cellular damage and cytokine storm.

## RESULTS

### Identification of novel CoV-RHIMs in SARS-CoV-2 Nsp13 and Nsp14 proteins

RHIMs consist of core tetrad (**`I/V-Q-I/V/L/C-G`**) residues promoting the stacking of β-sheet structures and formation of amyloid-like higher-order signaling complexes (*14, 15, 32, 39, 40*). Mutating these core tetrad residues inhibits the higher-order structure formation, cell death signaling, and inflammation (*15, 20, 21, 23, 32, 33, 41, 42*). We sought to identify the viral RHIM-like sequences in SARS-CoV-2 and other CoVs by protein sequence and structure-based analysis. Interestingly, we identified three novel RHIM-like sequences in Nsp13 (RNA helicase) and Nsp14 (exoribonuclease) of the SARS-CoV-2 and other CoVs **(Figure 1A, B)**. These RHIM-like CoV sequences (called CoV-RHIMs hereafter) were conserved in all the SARS-CoV-2 variants. The core tetrad in CoV-RHIM-1 and -3 of the SARS-CoV-2 showed high similarity with other viral and host RHIM-proteins, although the CoV-RHIM-2 was mutated in its core tetrad residues **(Figure 1B, 1D-F)**. In addition, a conserved Val/Ile (V/I) upstream of the core tetrad was noticeable in SARS-CoV-2 RHIM-proteins like established RHIM-proteins **(Figure 1B)**. More importantly, CoV-RHIMs of SARS-CoV-2 attained β-sheet conformation in crystal structures of Nsp13 and Nsp14 proteins and are surface accessible to be able to form complexes with other RHIM proteins when they are not in the replication complex (**Figure S1A**) (*43, 44*). These observations suggest that SARS-CoV-2 proteins show RHIM-like features and indicate their potential role in modulating host cell death and inflammatory responses.

**Figure 1:**
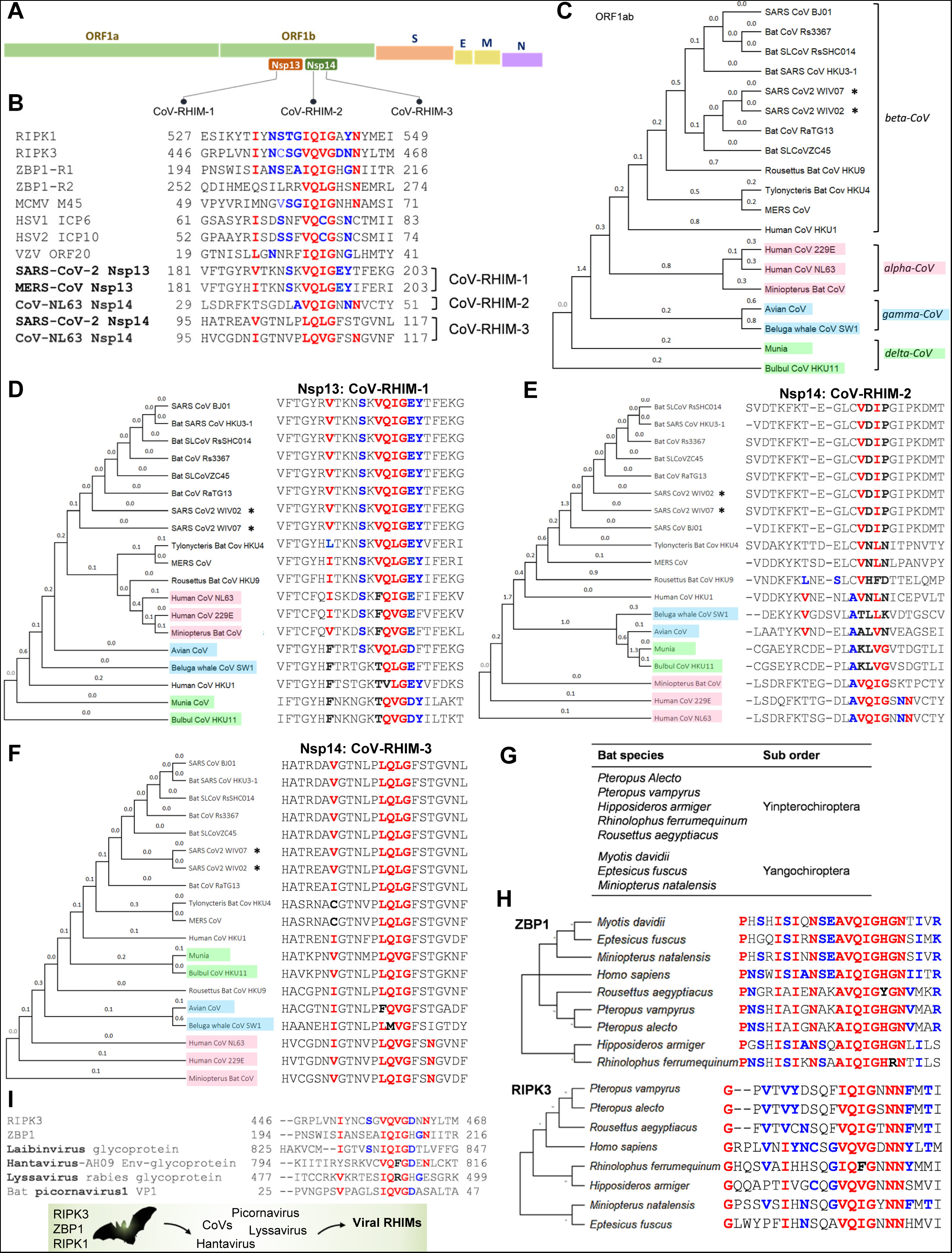
Identification of CoV-RHIMs in Nsp13 and Nsp14 of SARS-CoV-2 and bats as hosts in viral RHIM evolution in RNA viruses. **A,** Schematic representation showing the genomic organization of SARS-CoV-2 and CoV-RHIMs in Nsp13 and Nsp14 proteins encoded by ORF1b. **B,** Amino acid residues and their relative conservation in RHIMs of human proteins (RIPK1, RIPK3, and ZBP1), M45 of murine cytomegalovirus (MCMV), ICP6 and ICP10 of herpes simplex virus (HSV), ORF20 of varicella-zoster virus (VZV) and Nsp13 and Nsp14 of CoVs (SARS-CoV-2, MERS-CoV and CoV-NL63). Human ZBP1 consists of two RHIM sequences (ZBP1-R1 and ZBP1-R2). **C,** Phylogenetic tree representing the relatedness of different CoV genera based on protein sequences encoded by ORF1ab. **D-F,** The phylogenetic tree and the relative conservation of CoV-RHIM-1 in Nsp13 (**D**), CoV-RHIM-2 (**E**), and CoV-RHIM-3 (**F**) in Nsp14 across CoVs. **G,** Bat species from Yangochiroptera and Yinpterochiroptera suborders considered for analyzing bat RIPK3 and ZBP1 protein sequences. **H,** Protein sequence alignment and phylogenetic trees showing relative conservation of human and bat RHIM-sequences within RIPK3 and ZBP1 proteins. **I,** The potential viral RHIM signatures in lyssavirus rabies, picornavirus, and hantaviruses that originate from bats. The core tetrad residues and the proximal conserved residues of RHIMs are highlighted in bold; conserved or identical residue positions in RHIM sequences are highlighted in red; less conserved but chemically similar residue positions in RHIM sequences are highlighted in blue. Asterisk (*) symbols indicate Wuhan SARS-CoV-2 isolates.

Our further analysis of bat-originated beta-CoVs and other CoVs showed notable differences and distinct evolutionary patterns **(Figure 1C-F)** (*4, 45*). The CoV-RHIM-1 in Nsp13 was conserved in human-infected and bat beta-CoVs, including Bat-CoV RaTG13, which is one of the closed associates of SARS-CoV-2, suggesting the occurrence of CoV-RHIMs in bat-originated CoVs **(Figure 1D)**. HKU1 is a beta-CoV that infected humans after the SARS-CoV-1 outbreak but is associated with mild respiratory illness (*46*). HKU1 showed variation in the CoV-RHIM-1 and was distantly related to the pathogenic beta-CoVs in phylogenetic analysis **(Figure 1C and 1D)**. Human-CoV 229E and NL63 are alpha-CoVs causing mild respiratory symptoms like HKU1 (*46*). These CoVs also had a mutation at the first position of the core tetrad (V/I→F) **(Figure 1D)**. These observations further suggested an intriguing variation in CoV-RHIM-1 of the infected human CoVs that distinguishes mild and severe human CoV infections. We observed that avian-CoV, a gamma-CoV that did not infect humans so far, consisted of a conserved core tetrad, like pathogenic beta-CoVs **(Figure 1D)**. We identified two additional CoV-RHIMs in Nsp14 and Gln (Q) and Gly (G) at positions 2 and 4 of the core tetrad in CoV-RHIM-2 are mutated in pathogenic beta-CoVs **(Figure 1E and 1F)**. Unlike beta-CoVs, the alpha-CoVs (229E and NL63) consisted of the conserved residues promoting RHIM-mediated complex formation **(Figure 1E)**. Thus, the CoV-RHIMs show variations between alpha and beta-CoVs, which were zoonotically transmitted to humans but diverged in their pathogenicity. The CoV-RHIM-3 in Nsp14 is highly conserved across CoVs, except in gamma-CoVs. In addition, CoV-RHIM-3 of alpha-CoVs showed high similarity to human-RHIMs. These observations suggest that SARS-CoV-2 and other CoV proteins show RHIM-like features and indicate their potential role in modulating host cell death and inflammatory responses.

### Bats as the source of viral RHIM-evolution in RNA viruses

Why CoVs consist of multiple RHIM-like sequences in nonstructural proteins, and what factors drove the evolution of viral RHIMs in CoVs? Bats are the primary reservoirs of CoVs and identifying bat CoV-RaTG13 and other related viruses as the closest relative of the SARS-CoV-2 further elucidated the association of bat CoVs with severe human infections (*4, 6, 45, 47*). Also, bats are the natural reservoirs for other pathogenic RNA viruses that trigger cellular damage and inflammatory responses (*6, 47*). Bats are mammals with immune gene networks like humans (*48–50*). Bats also express RIPK3 and ZBP1, which are the potential targets for viral RHIM-mediated host modulation (*51, 52*). We observed that the core tetrad and surrounding residues in the RHIMs of bat RIPK3 and ZBP1 are conserved in both Yangochiroptera and Yinpterochiroptera suborders and are nearly identical to the RHIMs of human RIPK3 and ZBP1 (**Figure 1G,1H**). This raises the possibility that CoVs might have encountered the antiviral effects of bat RIPK3 and ZBP1 prior to their zoonotic transfer into humans. This suggests that zoonotic RNA viruses originating in bats might evolve to operate viral RHIMs. In line with this hypothesis, we found that lyssavirus rabies, picornavirus, and hantaviruses originated from bats encode RHIM-like sequences (**Figure 1I**). These observations thus suggest the likely evolution of viral RHIM-like sequences in bat RNA viruses. When the bat RNA viruses infect humans, they may either counteract or promote human RHIM-mediated cell death, contributing to virus propagation and severe pathology.

### CoV-RHIM-1 of 1B domain and RNA binding channel promote Nsp13-mediated cell death

To examine whether CoV-RHIMs are associated with cell death regulation in human cells, we generated lentiviruses expressing SARS-CoV-2 Nsp13-WT. We noticed robust cell death in HT-29 cells transduced with Nsp13-expressing lentiviruses but not EGFP-expressing lentiviruses, followed by puromycin selection (**Figure 2A**). The detection of Nsp13 protein expression in HT-29 cells indicated that the cell death was not due to defective Nsp13 lentiviruses (**Figure 2B**). A549 and HCT116 cells also showed robust cell death after Nsp13-lentivirus transduction (**Figure 2A,2B**). This suggests that SARS-CoV-2 Nsp13 expression promotes cell death in multiple human cell types. We also observed robust cell death in HT-29 cells transduced with SARS-CoV-1 and MERS-CoV Nsp13 lentiviruses (**Figure 2C**). SARS-CoV-2 Nsp13 consists of two RecA ATPase domains (Rec1A and Rec2A), which unwind DNA and RNA (*53, 54*). This helicase activity of Nsp13 is critical for the progressive elongation of the RNA-dependent RNA polymerase (RdRp) on a highly structured SARS-CoV-2 RNA genome (*53–55*). Nsp13 also consists of a Zinc-binding domain (ZBD), a stalk region (S), and an inserted domain 1B (hereafter called 1B) (**Figure 2D**). Rec1A, Rec2A, 1B, and a part of the S domain of the Nsp13 form an RNA binding channel, and the CoV-RHIM-1 spans to domain 1B (**Figure 2D**) (*53–55*). To understand how Nsp13 induces cell death, we generated strategic domain deletion constructs of the Nsp13, and immunoblotting analysis confirmed their expression at the protein level (**Figure S1B, S1C**). Deletion of 1B, Rec1A, and Rec2A or only Rec1A and Rec2A showed significant inhibition of Nsp13-mediated cell death (**Figure S1D**). This suggested the critical role of 1B, Rec1A, and Rec2A in Nsp13-dependent cell death. In addition, deletion of the ZBD or 1B domain did not diminish Nsp13-mediated cell death (**Figure S1D**). Interestingly, deletion of 1B and ZBD-S domains showed diminished Nsp13-mediated cell death (**Figure S1D**). These observations indicate that 1B domain deletion in combination with Rec1A-Rec2A or ZBD-S domains abolished Nsp13-dependent cell death. Thus, Nsp13-dependent cell death in human cells requires an RNA binding channel (formed by Rec1A, Rec2A, 1B, and a part of S) function.

**Figure 2:**
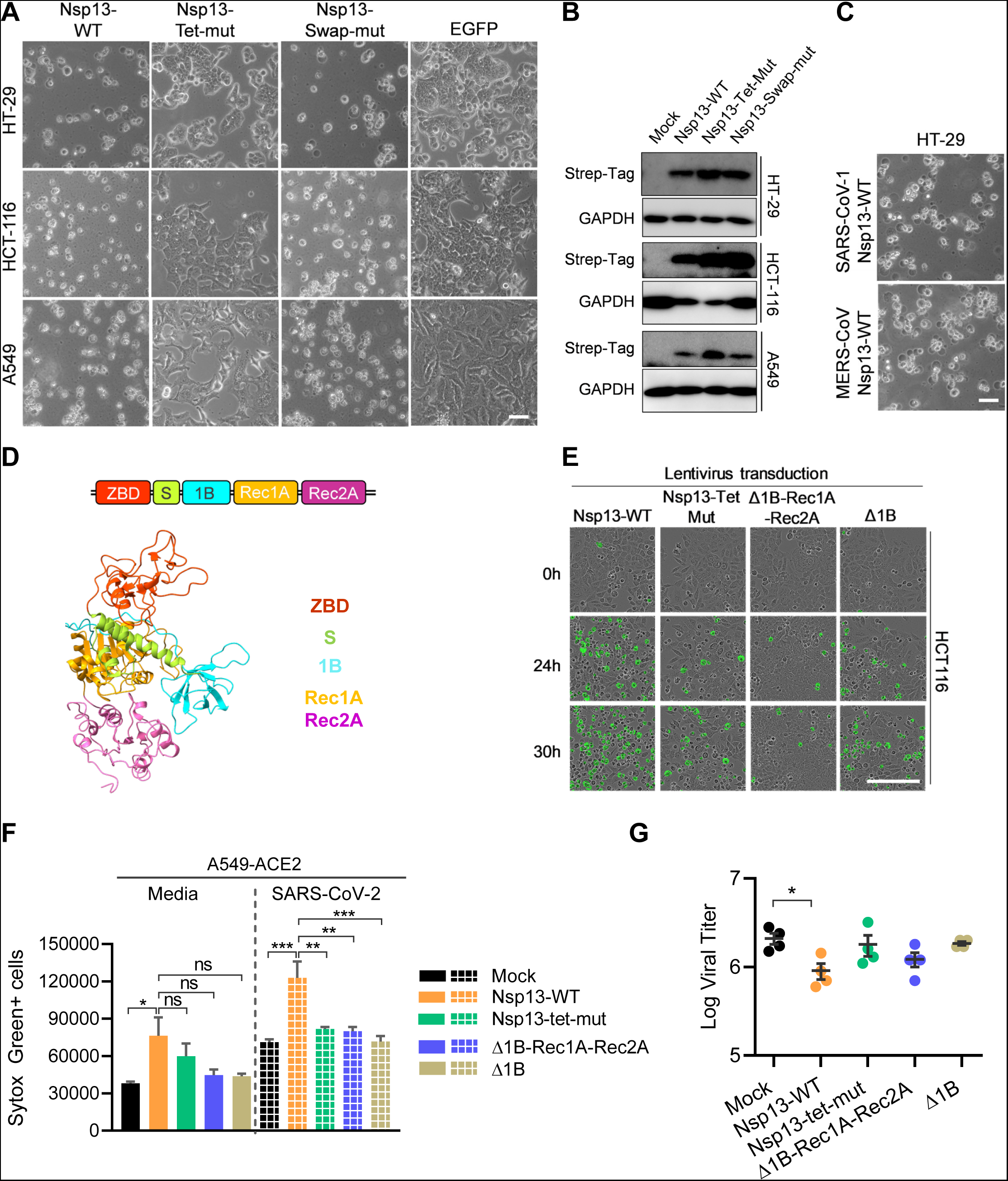
Nsp13 of pathogenic beta-CoVs promotes cell death in human cells and is dependent on CoV-RHIM-1 and RNA-binding channel. **A,** Microscopic analysis of HT-29, HCT-116, and A549 cells infected with lentiviruses expressing SARS-CoV-2 Nsp13-WT, Nsp13-Tet-mut, Nsp13-Swap-mut and EGFP followed by puromycin treatment. Scale bar, 50μm. **B,** Immunoblot analysis of lysates from HT-29, HCT-116, and A549 cells showing expression of Nsp13-WT, Nsp13-Tet-mut, Nsp13-Swap-mut (Strep-Tag) and GAPDH after lentivirus transduction. **C,** Microscopic analysis of HT-29 cells transduced with lentiviruses expressing SARS-CoV-1 and MERS-CoV Nsp13-WT. Scale bar, 50μm. **D,** The annotation of domains in the Nsp13 structure (PDB ID: 6ZSL). **E,** Representative Sytox green staining images of HCT-116 cells infected with lentiviruses expressing SARS-CoV-2 Nsp13-WT, Nsp13-Tet-mut, Nsp13-Δ1B and Nsp13-Δ1B-Rec1A-Rec2A acquired by Incucyte imaging analysis system. Scale bar, 200μm. **F,** Cell death levels in mock or SARS-CoV-2 infected A549-ACE2 cells ectopically expressing Nsp13 constructs as indicated. Cell death was monitored by Sytox green staining at 48 h of infection. \*\*\**P*=0.0002 (comparison of SARS-CoV-2 infected mock and Nsp13-WT cells), \*\*\**P*=0.0003 (comparison of SARS-CoV-2 infected Nsp13-WT and Δ1B expressing cells), \*\**P*=0.0076 (comparison of SARS-CoV-2 infected Nsp13-WT and Nsp13-Tet-mut expressing cells),\*\**P*=0.0043 (comparison of SARS-CoV-2 infected Nsp13-WT and Δ1B-Rec1A-Rec2A expressing cells), \**P*=0.0178, ns – not significant (one way ANOVA). **G,** SARS-CoV-2 infectivity titers in cell supernatants of mock or SARS-CoV-2 infected A549-ACE2 cells ectopically expressing Nsp13 constructs, measured by plaque assay. \**P*=0.0318 (one way ANOVA). Data shown are mean ± SEM.

To further understand the role of CoV-RHIM-1 in Nsp13-mediated cell death, we mutated the core tetrad residues of the CoV-RHIM-1 to alanine (VQIG→AAAA, named Nsp13-Tet-mut). We also swapped the CoV-RHIM-1 core tetrad of the SARS-CoV-2 Nsp13 with less pathogenic HKU1-CoV residues (VQIG→TVLG, named Nsp13-Swap-mut)). Nsp13-Tet-mut lentiviruses did not promote cell death in HT-29, A549, and HCT-116 cells, but Nsp13-Swap-mut induced cell death similarly to Nsp13-WT (**Figure 2A, 2B**). This implies that CoV-RHIM-1 is critical for Nsp13-mediated cell death in human cells. Nsp13-Swap-mut did not affect Nsp13-mediated cell death, perhaps due to conserved CoV-RHIM-1 conformation. Also, real-time monitoring of cell death using sytox green staining in HCT-116 cells transduced or transiently transfected with Nsp13 constructs showed reduced cell death when disrupting RNA binding channel or mutating CoV-RHIM-1 (**Figure 2E, S1E, S1G**). We also observed reduced apoptosis in HCT-116 cells when CoV-RHIM-1 was mutated or the RNA-binding channel was perturbed (**Figure S1F**). These observations indicate that CoV-RHIM-1 and RNA binding channel were required for Nsp13-mediated cell death. Perhaps RNA interaction with Nsp13 might facilitate RHIM-mediated cell death.

We then asked why mutating the CoV-RHIM-1 tetrad is sufficient to abolish Nsp13-mediated cell death, despite the requirement of 1B and RNA binding channel for cell death. The SARS-CoV-2 mini replication-transcription complex (RTC) consists of two Nsp13 molecules, one in apo-form (Nsp13-Apo) and the other in RNA-bound conformation (Nsp13-Rbound) (**Figure S2A**) (*53–55*). The two Nsp13 molecules are in proximity due to an interacting interface at domain 1B. Nsp13-Rbound shows a shift in its 1B domain orientation compared to NSP13-Apo, which is critical for facilitating fully open conformation of RNA binding channel for RNA entry (**Figure S2B-D**). Using AlphaFold, we predicted the structure of Nsp13-Tet-mut and found that it attains similar conformation as Nsp13-Apo and Nsp13-Rbound (**Figure S2E-G**). However, we observed striking conformational perturbations in the vicinity of the RNA binding channel of Nsp13-Tet-mut (**Figure S2B-D, S2E-G**). Structural superimposition indicated that Nsp13-Tet-mut differed from Nsp13-Apo in RNA binding channel conformation (**Figure S1E-G**). Nsp13-Tet-mut attained open RNA binding channel conformation similar to Nsp13-Rbound because of the significant shift in domain 1B orientation (**Figure S2G**). Furthermore, the Rec2A domain of the Nsp13-tet-mut showed conformational alterations compared to Nsp13-Rbound conformation. This implicated that tetrad mutations in CoV-RHIM-1 altered the conformation of domain 1B and Rec2A in the RNA binding channel of Nsp13, which explains the Nsp13-Tet-mut mediated resistance to cell death.

### Nsp13 promotes cell death in SARS-CoV-2 infected cells and is dependent on CoV-RHIM-1 and RNA binding channel

Mutations in Nsp13 are associated with species-specific and geographical adaptation of SARS-CoV-2, and targeting Nsp13 is known to impair SARS-CoV-2 replication (*56–59*). Also, substitutions in Nsp13 of CoVs severely affect replication and viral propagation (*58–62*). Thus, generating Nsp13 mutants of SARS-CoV-2 is challenging. To further understand the role of Nsp13 in SARS-CoV-2-induced cell death, we ectopically expressed Nsp13-WT, Nsp13-Tet-mut, and Nsp13 lacking RNA binding channel spanning domains (Δ1B-Rec1A-Rec2A) or only 1B domain (Δ1B) in A549 cells expressing human ACE2 (A549-ACE2). These cells were infected with SARS-CoV-2 (Hong Kong/VM20001061/2020), and cell death was measured in infected cells using Sytox green staining. Immunoblotting for SARS-CoV-2 nucleocapsid (N) protein in infected cells suggested productive SARS-CoV-2 infection and replication (**Figure S1H**). We found uninfected Nsp13-WT expressing A549-ACE2 cells showed increased basal-level cell death (**Figure 2F**). Upon SARS-CoV-2 infection, Nsp13-WT expressing A549-ACE2 cells showed significantly increased levels of cell death than infected mock cells (**Figure 2F**). However, Nsp13-Tet-mut, Δ1B-Rec1A-Rec2A, and Δ1B expressing cells showed diminished cell death than Nsp13-WT after SARS-CoV-2 infection (**Figure 2F**). Both uninfected and infected cells carrying Nsp13 showed increased levels of cell death compared to cells carrying Nsp13 CoV-RHIM-1 and RNA-binding channel mutants. This indicates that the ectopically expressed Nsp13 was dominant over virus-encoded Nsp13, consistent with overexpression studies, and suggested that Nsp13 promotes cell death in SARS-CoV-2 infected cells. To determine whether cell death is associated with SARS-CoV-2 virus propagation or curtail infection, viral titers in infected cell supernatants were measured using a Vero-E6 cell-based plaque assay. Nsp13-WT expressing cells showed lesser viral titers in supernatant compared to mock or Nsp13-Tet-mut, Δ1B-Rec1A-Rec2A, and Δ1B expressing cells upon SARS-CoV-2 infection (**Figure 2G**). Cell death rate and viral titers in SARS-CoV-2 infected Nsp13-WT expressing cells indicate that Nsp13-mediated cell death restricted SARS-CoV-2 viral propagation *in vitro*.

### CoV-RHIM-1 is less critical for Nsp13-mediated cell death in bat cells

Viral RHIMs are known to regulate species-specific cell death programs (*24, 31, 34*). Bats are the natural reservoir hosts for SARS-CoV-2-like viruses and likely promote viral RHIM evolution in them. Thus, we evaluated whether Nsp13 promotes or restricts RHIM-mediated cell death in a bat lung cell line (Tb1 Lu). Lentivirus virus-based SARS-CoV-2 Nsp13 expression promoted cell death in Tb1 Lu cells like in human cells (**Figure 3A, 3B**). Sytox green staining indicated a lower magnitude of cell death than human cells (**Figure 3C, 3D**). However, Caspase-3/7 dye and Sytox green staining indicated that Nsp13 expressing Tb1 Lu cells showed a high magnitude of apoptosis and detectable levels of necrotic cell death. Interestingly, mutating CoV-RHIM-1 in Nsp13 significantly decreased apoptosis (Caspase-3/7) compared to necrotic cell death (Sytox green) (**Figure 3C**, 3D). Domain deletions affecting RNA binding channel diminished apoptosis and necrotic cell death in Nsp13 expressing Tb1 Lu cells. Also, transient expression of Nsp13 induced Tb1 Lu cell death in a dose-dependent manner and disrupting RNA binding channel reduced cell death than CoV-RHIM-1 mutant (**Figure S3A-D**). These observations indicate that Nsp13 promoted cell death, preferentially apoptosis, in bat cells. Loss of RNA binding channel function diminished Nsp13-mediated cell death, whereas mutating CoV-RHIM-1 in Nsp13 marginally alleviated Nsp13-mediated cell death, unlike in human cells.

**Figure 3:**
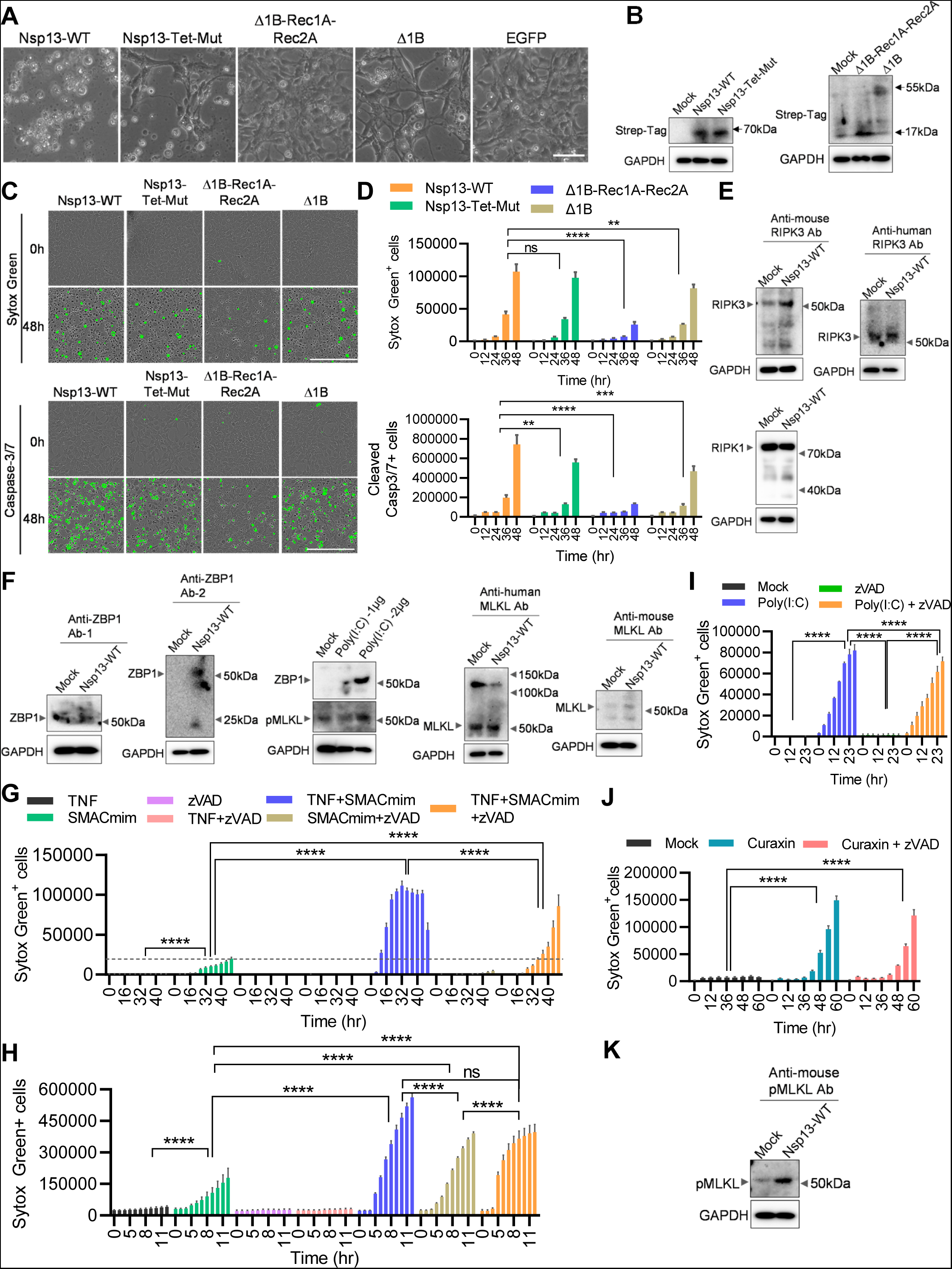
Nsp13 promotes RNA-binding channel dependent bat cell death and CoV-RHIM-1 function is less critical. **A,** Microscopic analysis of Tb1 Lu cells infected with lentiviruses expressing SARS-CoV-2 Nsp13-WT, Nsp13-Tet-mut, Nsp13-Δ1B, Nsp13- Δ1B-Rec1A-Rec2A and EGFP followed by puromycin treatment. Scale bar, 50μm. **B,** Immunoblot analysis of lysates from Tb1 Lu cells showing expression of Nsp13-WT and its mutants after lentivirus transduction. **C,** Representative Caspase-3/7 and Sytox green staining images of Tb1 Lu cells infected with lentiviruses expressing SARS-CoV-2 Nsp13-WT, Nsp13-Tet-mut, Nsp13-Δ1B and Nsp13-Δ1B-Rec1A-Rec2A followed by puromycin treatment acquired by Incucyte imaging analysis system. Scale bar, 200μm. **D,** Real-time cell death measurement by Caspase-3/7 and Sytox green staining of Tb1 Lu cells infected as in panel-C. \*\*\*\**P*<0.0001, \*\*\**P*=0.0003, \*\**P*=0.0046 (Sytox green staining), \*\**P*=0.0086 (Caspase-3/7 staining) (two way ANOVA, n=3). Data shown are mean ± SEM. **E & F**, Immunoblot analysis of RIPK1, RIPK3, ZBP1 and MLKL in lysates from ‘Tb1Lu cells after mock treatment, Nsp13 or Poly(I:C) transfection. The antibodies used for detecting these proteins were specific to human and mouse proteins. **G & H,** Real-time cell death measurement by Sytox green staining of Tb1 Lu (**G**) and HT-29 cells (**H**), after treatment of TNF, zVAD and SMACmimetic (SMACmim). \*\*\*\**P*<0.0001 (two way ANOVA, n=3), ns – not significant (two way ANOVA). **I & J,** Real-time cell death measurement by Sytox green staining of Tb1 Lu cells transfected with Poly(I:C) **(I)** or treated with Curaxin **(J)** \*\*\*\**P*<0.0001 (two way ANOVA, n=3). Data shown are mean ± SEM. **K**, Immunoblot analysis of phosphorylated MLKL (pMLKL) in lysates from Tb1Lu cells after mock or Nsp13 transfection.

The expression of RHIM-proteins and activation of RHIM-mediated apoptosis and necroptosis in bat cells is unknown. We tested the expression of RHIM-proteins in Tb1 Lu cells using multiple commercially available antibodies against RIPK1, RIPK3, and ZBP1. We found detectable protein levels of RIPK1, RIPK3, and ZBP1 in Tb1 Lu cells (**Figure 3E, 3F**). Transfection of double-stranded RNA ligand, Poly(I:C), upregulated ZBP1 expression in Tb1 Lu cells (**Figure 3F**). We also observed the MLKL expression in Tb1 Lu cells (**Figure 3F**). Notably, we observed lower expression levels of RHIM proteins and MLKL in bat cells compared to human cells. This discrepancy in detection levels might be due to the antibodies specific to detect human and mouse proteins. The detectable levels of RHIM-proteins led us to test whether bat cells undergo RHIM-mediated apoptosis and necroptosis. TNF, zVAD or TNF + zVAD treatment did not induce cell death, however, TNF or TNF + zVAD in combination with SMACmimetic (SMACmim) induced robust Tb1 Lu cell death (**Figure 3G**). However, the cell death kinetics were slower than human cells (**Figure 3G, 3H**). Adding GW806742X, an MLKL-specific inhibitor, diminished TNF-induced necroptosis activation in Tb1 Lu and HT-29 cells (**Figure S3E, S3F**). GW806742X mediated necroptosis inhibition was more robust in HT-29 cells than Tb1 Lu cells. Poly(I:C) or ZBP1 activating Curaxin (CBL0137) also induced Tb1 Lu cell death (**Figure 3I, 3J**). This suggests the sensitivity of Tb1 Lu cells for RHIM-protein-mediated apoptosis and necroptosis. In line with these observations, we observed detectable MLKL phosphorylation in Tb1 Lu cells with an antibody specific to mouse MLKL phosphorylated at Ser345 (Ser358 in humans), suggesting MLKL activation in those cells during necroptosis activation (**Figure S3G**). These observations suggest that bat Tb1 Lu cells undergo cell death in response to RHIM-dependent apoptosis and necroptosis triggers. Also, ectopic expression of Nsp13 in Tb1 Lu cells promoted phosphorylation of MLKL, suggesting host RHIM-protein mediated signalign activation by Nsp13 (**Figure 3K**). These observations in a bat cell line suggest that Nsp13 promotes cell death in bat cells like human cells. However, unlike human cells, the bat cell death mainly depends on the RNA binding channel than the CoV-RHIM-1, indicating host-specific Nsp13 mediated cell death regulation. It is unknown why bat cells preferentially activate RHIM-dependent apoptosis, unlike human cells. Also, to the best of our knowledge, we show for the first time that bat cells operate RHIM-dependent cell death programs.

### SARS-CoV-2 Nsp13 shows RHIM-dependent association with human RHIM-proteins

RHIM-RHIM interactions are critical for the assembly of cell death signaling complexes like necrosome and ripoptosome to activate apoptosis and necroptosis (*13, 14, 63, 64*). To understand Nsp13 CoV-RHIM-1 role in host-RHIM-driven cell death, we ectopically expressed Nsp13 and its mutants in HT-29 cells and treated them with TNF and SMACmimetic (TNF+SMACmim) for TNF-induced apoptosis and TNF+SMACmim+zVAD for TNF-induced necroptosis (**Figure S4A**). As expected, these triggers induced robust cell death in HT-29 cells (**Figure S4A**). Expression of Nsp13-WT, Nsp13-Tet-mut, or Nsp13-Swap-mut did not diminish TNF-induced apoptosis and necroptosis in HT-29 cells, suggesting a dispensable role of Nsp13 in RIPK1-RIPK3 signaling (**Figure S4A**). RHIM-mediated RIPK1 and RIPK3 interactions are critical for TNF-induced apoptosis and necroptosis (*15, 17, 18, 42, 65*). Viral RHIM-proteins, M45, and ICPs directly interact with RIPK3 and ZBP1 through RHIM-homotypic interactions and restrict virus-induced necroptosis (*11, 20, 21, 24*). ORF20 of VZV also interacts with ZBP1 to regulate virus-induced apoptosis (*33*). HSV-1 viral RHIM-protein, ICP6, shows species-specific necroptosis regulation by promoting viral-RHIM mediated necroptosis in mouse cells but restricting necroptosis activation in human cells (*24, 31, 34*). We sought to determine the ability of SARS-CoV-2 Nsp13 to interact with human RIPK1, RIPK3, and ZBP1 by co-expressing these proteins in HEK-293T cells, followed by immunoprecipitation. HEK-293T cells express RIPK1 endogenously but do not express RIPK3 and ZBP1 proteins. When Nsp13, RIPK3, and ZBP1 were co-expressed in HEK-293T cells, immunoprecipitation with anti-HA-tag (RIPK3-HA) and anti-ZBP1 antibodies showed RIPK3-RIPK1-ZBP1 complex formation without Nsp13 (**Figure 4A, 4B**). Nsp13 did not show detectable interaction with RIPK1 and RIPK3, corroborating the dispensable role of Nsp13 in TNF-induced apoptosis and necroptosis (**Figure 4A, 4B**). However, Nsp13 immunoprecipitation showed its interaction with ZBP1. This suggests that Nsp13 can interact with ZBP1 but may not interfere with host RHIM-protein interactions. To understand whether these interactions are RHIM-dependent, we sought to test the interaction of RHIM and 1B domain deletion mutants of Nsp13 with the ZBP1 through immunoprecipitation. Nsp13-WT interacted with the ZBP1-WT, but Nsp13-Tet-mut, and the Nsp13 lacking 1B did not interact with the ZBP1-WT (**Figure 4C, 4D**). Also, mutating the RHIM domain (RHIM-1) of the ZBP1 disrupted its interaction with the Nsp13-WT (**Figure 4C, 4D**). These observations further indicate that the association of Nsp13 with the ZBP1 is RHIM-dependent.

**Figure 4:**
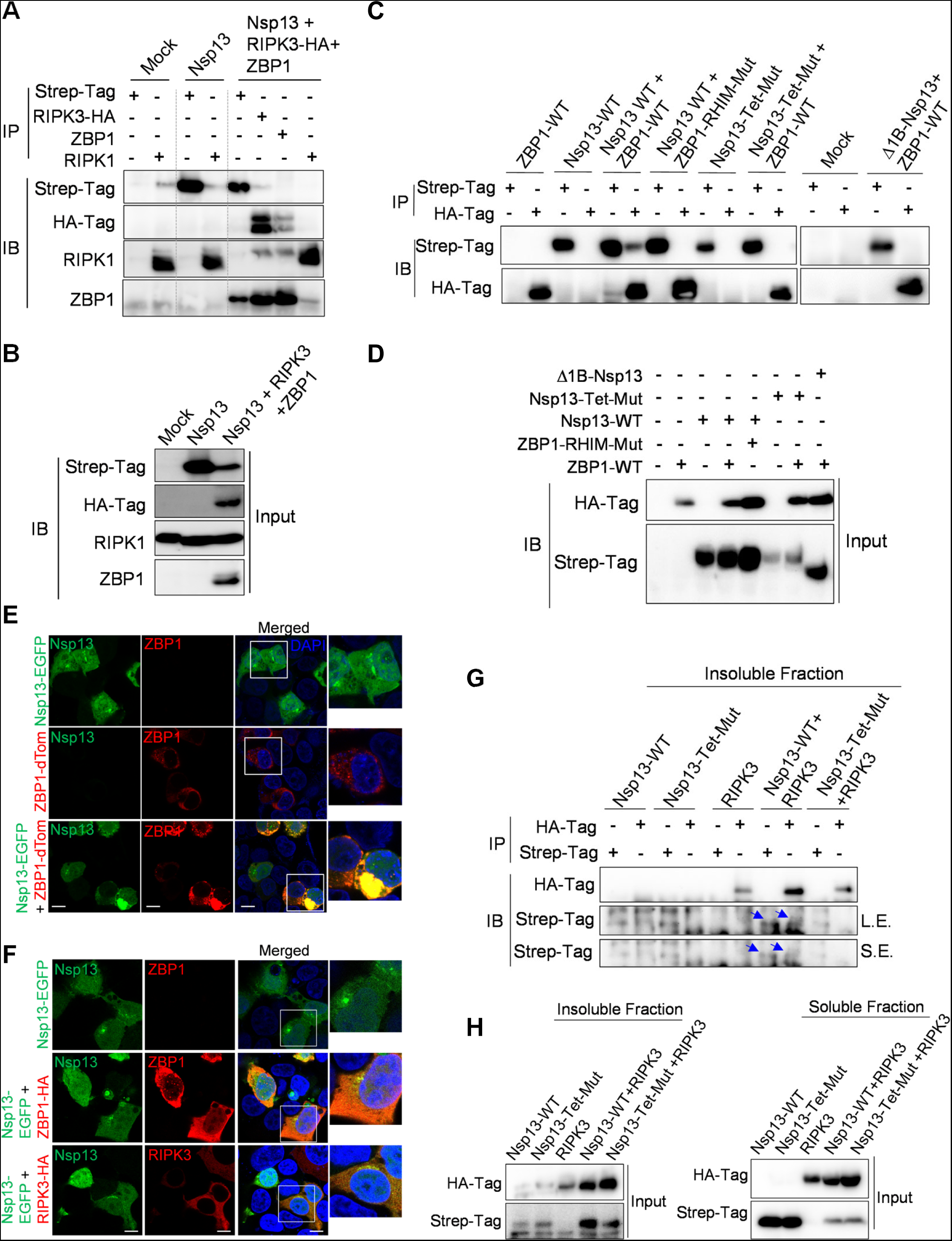
SARS-CoV-2 Nsp13 show RHIM-dependent interaction with host ZBP1 and RIPK3 proteins. **A-B,** Immunoblot analysis of anti-Strep-Tag (Nsp13), anti-RIPK1, anti-HA-tag (RIPK3), and anti-ZBP1 immunoprecipitates **(A)** and whole cell lysates (inputs) **(B)** from HEK-293T cell lysates expressing Nsp13 alone or co-expressing Nsp13, RIPK3 and ZBP1. **C-D,** Immunoblot analysis of anti-Strep-Tag (Nsp13) and anti-HA-tag (ZBP1) immunoprecipitates **(C)** and inputs **(D)** from HEK-293T cell lysates expressing Nsp13 WT, Nsp13-Tet-Mut, Δ1B, ZBP1-WT and ZBP1-RHIM-Mut alone or in indicated combinations of Nsp13 and ZBP1 constructs. **E,** Confocal microscopy imaging of HEK-293T cells expressing Nsp13-EGFP or ZBP1-dTomoto alone or co-expressing both the constructs. Scale bar, 10μm. **F,** Confocal microscopy imaging of HEK-293Tcells expressing Nsp13-EGFP and in combination with ZBP1-HA or RIPK3-HA. Scale bar, 10μm.**G,** Immunoblot analysis of anti-Strep-Tag (Nsp13) and anti-HA-tag (RIPK3) immunoprecipitates from insoluble fractions of HEK-293T cell lysates expressing Nsp13-WT, Nsp13-Tet-Mut and RIPK3 individually or in indicated combinations of Nsp13 and RIPK3 constructs. **H,** Inputs for insoluble and soluble fractions of cell lysates for Panel-G.

To further establish SARS-CoV-2 Nsp13 association with host-RHIM proteins, we tagged Nsp13 with EGFP (Nsp13-EGFP) for capturing Nsp13 association with RHIM proteins in imaging studies (**Figure S4B-D**). Tagging EGFP did not alter Nsp13-mediated cell death and protein expression, indicating the retention of Nsp13 native conformation upon EGFP fusion (**Figure S4C, S4D**). Nsp13-EGFP and ZBP1-dTomato were colocalized upon co-expression, and the colocalized regions appeared as complexes (**Figure 4E**). We further co-expressed Nsp13-EGFP with ZBP1 tagged with HA-Tag (ZBP1-HA) and found that Nsp13 and ZBP1 colocalize in HEK-293T cells (**Figure 4F**). Also, Nsp13 appeared to be colocalized with RIPK3 but not RIPK1 (**Figure 4F, S4E**). Real-time analysis of Nsp13-EGFP and ZBP1-dTomato expression in live cells showed colocalization of Nsp13 with the ZBP1 upon expression and led to cell death phenotype (**Figure S4F**). The co-expression of Nsp13 with the RHIM-mutant of ZBP1 reduced Nsp13-ZBP1 colocalization, suggesting RHIM-dependent association (**Figure S4G, S4H**). These observations further indicate the association of Nsp13 with the ZBP1 and RIPK3 and higher-order complex formation. Although we did not observe Nsp13 and RIPK3 interaction in immunoprecipitation experiments, imaging studies show that Nsp13 colocalized with RIPK3 (**Figure 4F**). To validate this further, we separated soluble and insoluble fractions from the whole cell lysates co-expressing Nsp13-WT or Nsp13-Tet-mut with RIPK3 and subjected to immunoprecipitation with Strep-tag and HA-tag antibodies. We observed detectable interaction of RIPK3 with Nsp13 in the insoluble fraction but not in the soluble fraction (**Figure 4G, S4I**). Moreover, Nsp13 and RIPK3 were present in soluble fractions when expressed individually, and significant protein fractions of Nsp13 and RIPK3 appeared in insoluble fractions when co-expressed (**Figure 4H**). These observations further indicate the association of Nsp13 with RIPK3 and ZBP1.

### SARS-CoV-2 Nsp13 promotes the formation of large insoluble complexes of RHIM proteins

RHIM-RHIM interactions of host and viral proteins form fibrillar insoluble oligomeric complexes (*14, 32, 63, 66*). Recent studies demonstrated the formation of insoluble amyloid-like complexes by RIPK1, RIPK3 and ZBP1 upon interaction with viral-RHIM proteins (*14, 32, 63, 66*). To monitor whether SARS-CoV-2 Nsp13 can assemble amyloid-like complexes, we expressed Nsp13 and its mutants alone or in combination with RIPK3 and ZBP1 in HEK-293T cells. To preserve oligomeric complexes, cell lysates were crosslinked using a DSP cross-linker, which has a built-in disulfide in its spacer region that allows decoupling of crosslinked oligomers through treatment with disulfide reducing agents such as beta-mercaptoethanol (BME). Nsp13-WT expression led to the formation of NP-40 soluble higher-order complexes with undetectable oligomers in the insoluble fraction (**Figure S5A**). We observed an increased complex formation by Nsp13-Tet-mut and Nsp13-Swap-mut than Nsp13-WT in NP40-soluble fractions (**Figure S5A**). Nsp13-Swap-mut also formed detectable NP-40 insoluble oligomers. Thus, SARS-CoV-2 Nsp13 appeared to oligomerize into large complexes upon overexpression and mutating or swapping core tetrad of CoV-RHIM-1 further enhanced homo-oligomer formation. This indicated the predominant monomeric nature of the Nsp13-WT protein, which might oligomerize when 1B domain conformation is perturbed. Since Nsp13 interacted with ZBP1 in HEK-293T cells, we further monitored whether Nsp13 affects ZBP1 oligomerization. We observed detectable but low levels of higher-order oligomers of ZBP1 in NP40-insoluble fraction upon transfection (**Figure S5A**). Co-expression of ZBP1 with Nsp13-WT showed significantly increased levels of ZBP1 oligomers in NP-40 insoluble fraction (**Figure S5A**). However, Nsp13-Tet-mut and ZBP1 co-expression showed a lesser magnitude of ZBP1 oligomer formation (**Figure S5A**). In contrast to the predominant monomeric form of Nsp13-WT, its co-expression with ZBP1 showed detectable levels of Nsp13-WT complexes in NP40-insoluble fraction and increased levels of oligomers in NP40-soluble fraction (**Figure S5A**). These crosslinking experiments suggested that Nsp13 promotes CoV-RHIM-dependent higher-order complexes of ZBP1. As Nsp13-Tet-mut already forms non-native oligomers, its expression with ZBP1 did not facilitate ZBP1 oligomerization (**Figure S5A**).

We further co-expressed Nsp13-WT with RIPK3 and ZBP1 and monitored higher-order complex formation. The co-expression of ZBP1 and RIPK3 led to the formation of NP-40 soluble and insoluble RIPK3 oligomers (**Figure 5A**). These oligomers were dissolved into monomers upon BME treatment (**Figure 5A**). However, ZBP1 did not show detectable oligomerization in both fractions (**Figure 5A**). Similarly, the co-expression of Nsp13-WT with RIPK3 triggered RIPK3 complex formation in NP40-insoluble and soluble fractions. Higher order complexes of the RIPK3 were observed even when RIPK3 was expressed with Nsp13-WT and ZBP1. The magnitude of oligomer formation was lesser than that was seen in Nsp13-WT and RIPK3 co-expressed cell lysates (**Figure 5A**). ZBP1 formed NP40-insoluble complexes when co-expressed with Nsp13-WT or Nsp13-WT and RIPK3 (**Figure 5A**). Even though BME treatment decoupled the complexes of RIPK3 and ZBP1, a fraction of complexes was not completely dissolved after BME treatment, suggesting the formation of insoluble fibrillar amyloid-like oligomers with hydrophobic interfaces as reported previously for host RHIM-proteins (**Figure 5A**) (*32, 63*). These observations suggest that Nsp13 forms soluble higher-order complexes when co-expressed with RIPK3 or ZBP1. Nsp13 co-expression promotes insoluble amyloid-like complexes of RIPK3 and ZBP1. Thus, Nsp13 may preferably form soluble oligomers that further nucleate RIPK3 and ZBP1 insoluble fibrillar amyloid-like necrosome assemblies.

**Figure 5:**
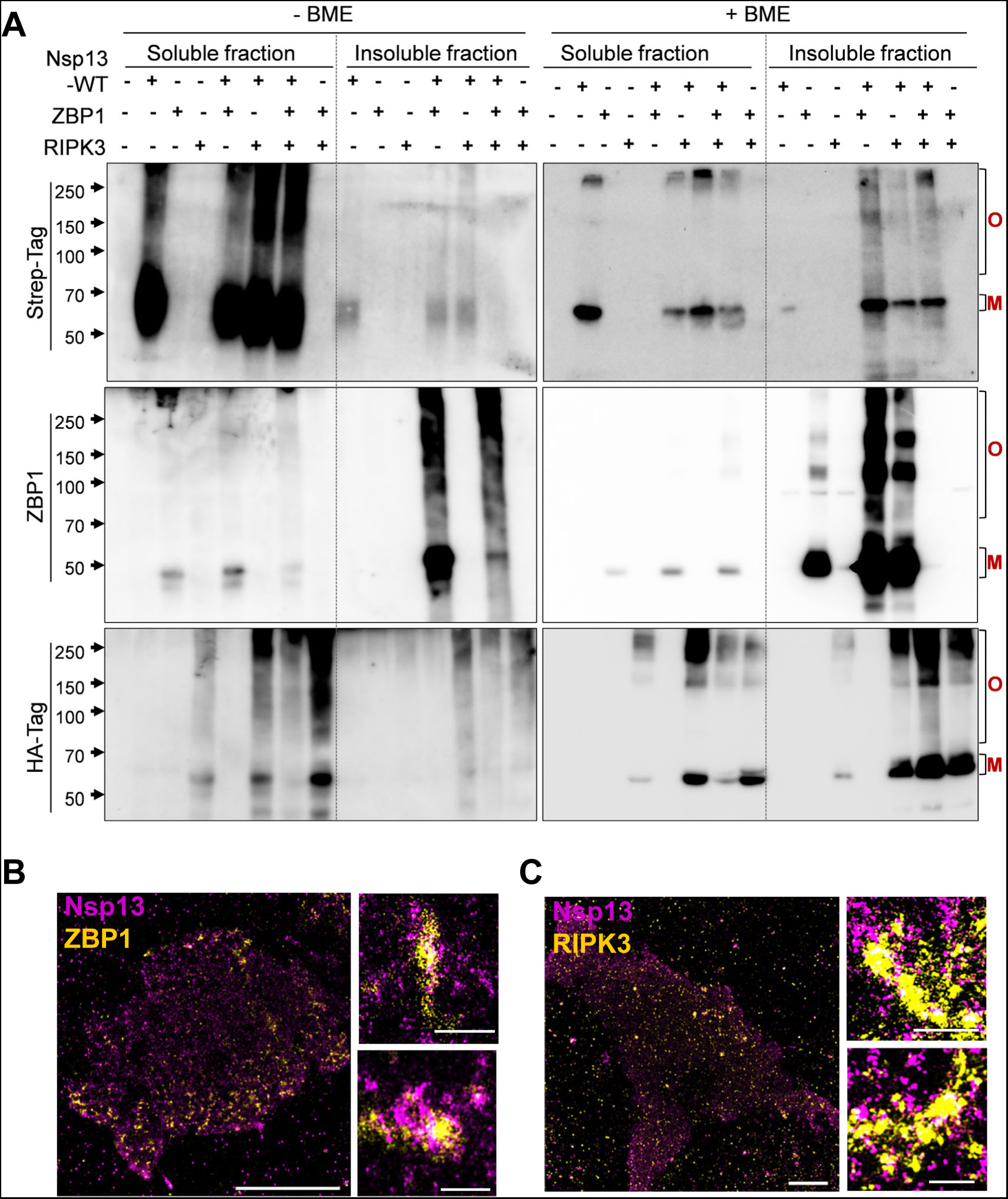
SARS-CoV-2 Nsp13 triggers host-RHIM protein oligomerization and large complex formation. **A,** Immunoblot analysis of crosslinked lysates of HEK-293T cells expressing Nsp13-WT, RIPK3 and ZBP1 individually or co-expressing Nsp13-WT+ZBP1, Nsp13-WT+RIPK3, Nsp13-WT+ZBP1+RIPK3 or ZBP1+RIPK3, in non-reduced (without BME) and reduced (with BME) conditions. O-oligomer complexes; M-Monomer. **B-C,** Visualization of SARS-CoV-2 Nsp13 and ZBP1 (**B**) or RIPK3 (**C**) in HEK-293T cells using DNA-PAINT imaging. Scale bars, 5 μm (complete cell image); 200nm (magnified images).

To further understand the association of Nsp13 with ZBP1 and RIPK3, we visualized Nsp13, ZBP1, and RIPK3 in HEK-293T cells using DNA-PAINT imaging that enables nanometer-scale resolution for monitoring complex formation. DNA-PAINT imaging suggested spatial localization of Nsp13 with ZBP1 and RIPK3 (**Figure 5B, 5C, S5B, S5C**). A recent study showed the formation of round and rod-shaped RHIM-protein amyloid complexes using super-resolution microscopy (*66*). ZBP1 appeared as rod-shaped complexes in Nsp13-ZBP1 complexes compared to Nsp13 (**Figure 5B, S5B**). RIPK3 appeared as round and rod-shaped structures in Nsp13-RIPK3 complexes (**Figure 5C, S5C**). The DNA-PAINT imaging indicated the formation of round and rod-shaped complexes by ZBP1 and RIPK3. Nsp13 is in proximity with ZBP1 and RIPK3 in complexes, although Nsp13 appeared to form smaller and more distinct structures than ZBP1 and RIPK3. Together, these observations indicate that Nsp13 employs viral RHIMs to promote the higher-order complex formation.

### SARS-CoV-2 Nsp13 promotes ZBP1-RIPK3 signaling and cell death

Higher-order complex formation by ZBP1 and Nsp13 after co-expression led us to examine the role of Nsp13 in ZBP1-RIPK3 signaling-specific cell death activation. ZBP1 activation triggers IAV-induced necroptosis, apoptosis and pyroptosis cell death programs. Most transformed human cell lines do not express RIPK3 and ZBP1. L929 cells (mouse fibroblast cell line) retain ZBP1 expression and show ZBP1-dependent cell death after IAV infection. Thus, we generated *Zbp1*^-/-^ L929 cells to study the role of Nsp13 in ZBP1-dependent cell death pathways. We transiently expressed Nsp13 in WT and *Zbp1*^-/-^ L929 cells and infected these cells with IAV to study the role of Nsp13 in IAV-induced necroptosis (**Figure 6A**). Without Nsp13 expression, *Zbp1*^-/-^ cells showed significantly less cell death than WT cells after IAV infection or IAV combined with zVAD treatment (IAV + zVAD, for activating necroptosis). Nsp13-WT expression in WT cells did not alter IAV or IAV+zVAD-induced cell death and promoted faster kinetics of cell death activation (**Figure 6A**). Interestingly, NSP13-WT expressing *Zbp1*^-/-^ cells showed a higher magnitude of cell death, which is comparable to WT-cells, suggesting the role of Nsp13 in promoting IAV-induced cell death in the absence of ZBP1 (**Figure 6A**). Immunoblotting analysis showed that Nsp13-WT expression in *Zbp1*^-/-^ cells led to increased pMLKL levels than *Zbp1*^-/-^ cells without Nsp13 expression (**Figure S5D**). This indicated IAV+zVAD-mediated necroptosis activation by Nsp13 despite lacking ZBP1 expression. Also, we observed increased cleaved caspase-3 in Nsp13-WT expressing *Zbp1*^-/-^ cells and no apparent differences in pMLKL levels (**Figure S5D**). However, Nsp13-Tet-mut expression in *Zbp1*^-/-^ cells did not reduce IAV or IAV+zVAD-induced cell death, suggesting that CoV-RHIM-1 mutations do not appear to abolish Nsp13-induced cell death in IAV-infected *Zbp1*^-/-^ cells (**Figure 6A**). Nsp13-WT and Nsp13-Tet- mut expressing WT and *Zbp1*^-/-^ cells showed a very low magnitude of cell death without infection than IAV and IAV+zVAD treatment (**Figure 6A**). To further understand whether Nsp13 promotes ZBP1-dependent cell death in human cells, we infected HT-29 cells with IAV+zVAD. HT-29 cells express RIPK3 but show undetectable ZBP1 expression. Expression of human ZBP1 in HT-29 cells led to increased necroptosis activation after IAV+zVAD treatment (**Figure 6B**). Nsp13-WT co-expression with ZBP1 enhanced IAV+zVAD-induced necroptosis in HT-29 cells (**Figure 6B**). Thus, it appeared that Nsp13 promoted ZBP1-induced cell death during IAV infection. Intriguingly, Nsp13 promoted IAV-induced cell death in the absence of ZBP1 expression.

**Figure 6:**
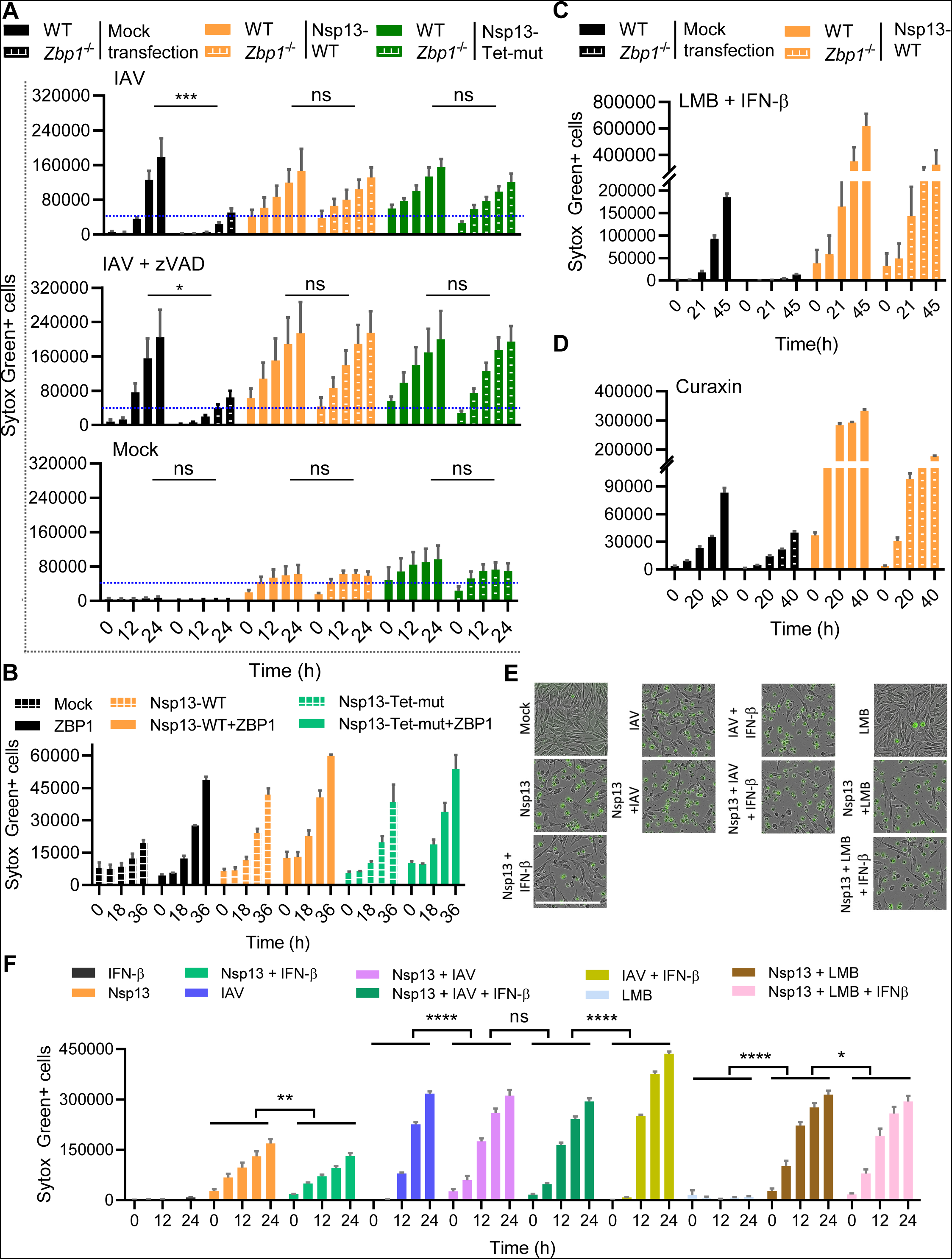
SARS-CoV-2 Nsp13 promotes ZBP1-RIPK3 signaling-dependent cell death and is regulated by intracellular RNA ligands. **A,** Real-time cell death measurement by Sytox green staining of WT and *Zbp1*^-/-^ L929 cells ectopically expressing Nsp13-WT and Nsp13-Tet-mut after IAV, IAV+zVAD and mock infection. \*\*\**P*=0.0005, \**P*=0.0307 (two-way ANOVA). **B,** Real-time cell death measurement by Sytox green staining of ZBP1 expressing HT-29 cells after Nsp13 transfection and IAV+zVAD infection. **C & D,** Cell death measurement by Sytox green staining of WT and *Zbp1*^-/-^ L929 cells expressing Nsp13-WT after LMB+IFN-β (**C**) and Curaxin (**D**) treatment. Data shown are mean ± SEM. **E,** Microscopic images of cell death in L929 treated as indicated and acquired by Incucyte imaging analysis for the cells. Scale bar, 200μm. **F,** Real-time analysis of cell death of mock or Nsp13 transfected L929 cells treated /infected with IFN-β alone, IAV alone, IAV + IFN-β, LMB alone or LMB + IFN-β. \*\*\*\**P*<0.0001, \*\**P*=0.0022, \**P*=0.0310 (two way ANOVA, *n*=3). Data shown are mean ± SEM.

Leptomycin-B (LMB), in combination with IFNs or curaxin (CBL0137) treatment, is known to trigger ZBP1 activation and cell death (*41, 67*). We examined LMB+IFN-β and curaxin-mediated cell death in WT and *Zbp1*^-/-^ L929 cells to further probe the role of SARS-CoV-2 Nsp13 in ZBP1 function (**Figure 6C,6D**). As expected, LMB+IFN-β treatment triggered cell death and lack of ZBP1 significantly abolished this cell death (**Figure 6C**). However, upon Nsp13-WT expression in *Zbp1*^-/-^ cells, increased cell death was observed after LMB+IFN-β treatment compared to those without Nsp13-WT expression (**Figure 6C**). Curaxin treatment in WT and *Zbp1*^-/-^ cells showed ZBP1-dependent cell death activation, and Nsp13-WT expression led to increased cell death in *Zbp1*^-/-^ cells similar to that of LMB+IFN-β treatment (**Figure 6D**). Nsp13-WT overexpression further enhanced cell death in WT cells after LMB+IFN-β or curaxin treatment (**Figure 6C, 6D**). These results further suggest that SARS-CoV-2 Nsp13 promotes ZBP1-mediated cell death, and in the absence of ZBP1 expression, Nsp13 triggers cell death in response to ZBP1 activating ligands. Why might Nsp13 have evolved to regulate ZBP1-RIPK3 signaling and induce host cell death? Type I interferons (IFN) promote ZBP1 and RIPK3 signaling and activation of cell death in viral infections (*9, 22, 41, 42, 68*). Delayed type I IFN production and restricted IFN signaling due to Nsp13 and other proteins of SARS-CoV-2 facilitate host modulation and efficient virus propagation (*1, 8*). We anticipate that the dampened type I IFN signaling might restrict early activation of ZBP1-RIPK3 signaling and cell death, which preferentially destroy viral replication platforms. Perhaps Nsp13 has evolved to trigger cell death due to insufficient ZBP1-RIPK3 signaling activation at the late stages of the infection to facilitate viral spread, tissue damage, and inflammation. Indeed, recent studies demonstrate IFN and inflammatory cytokine-driven cell death, tissue damage, and disease pathogenesis during SARS-CoV-2 infection at later stages (*1, 12, 69–71*).

### Intracellular RNA ligands and type I IFNs regulate SARS-CoV-2 Nsp13-mediated cell death

Type I IFNs upregulate ZBP1 and MLKL expression and promote necroptosis and other programmed cell death activation in viral infections (*9, 41, 42, 68*). SARS-CoV-2 Nsp13 is associated with innate immune evasion and restricts type I IFN signaling activation (*8*). We hypothesize that type I IFN treatment may regulate Nsp13-mediated cell death. Ectopic expression of Nsp13 in L929 cells promoted spontaneous cell death at the basal level, and IFN-β treatment partly reduced this Nsp13-induced cell death, suggesting IFN receptor-mediated cell death regulation (**Figure S6A, S6B**). Nsp13-mediated cell death requires CoV-RHIM in 1B domain and RNA binding channel (spanning 1B, Rec1A and Rec2A domains), suggesting RNA binding as a critical step for cell death activation by Nsp13. Intracellular delivery of Poly(I:C) (transfection) increased Nsp13-mediated cell death, but the addition of Poly(I:C) to the cell surface did not alter Nsp13-mediated cell death (**Figure S6C, S6D**). As expected, IAV infection in Nsp13- expressing cells promoted increased cell death (**Figure 6E, 6F**). However, unlike IFN-β- dependent inhibition of Nsp13-mediated cell death, IFN-β treatment after IAV infection in Nsp13- expressing cells did not reduce cell death (**Figure 6E, 6F**). Without Nsp13 expression, IAV and IFN-β treatment showed a further increase in cell death levels. Furthermore, LMB treatment in Nsp13-expressing cells showed a higher magnitude of cell death activation than in Nsp13- expressing untreated cells (**Figure 6E, 6F**). LMB+IFN-β treatment did not alter Nsp13-mediated cell death. These observations suggested that IFN-β did not inhibit Nsp13-mediated cell death during IAV infection or LMB treatment, which generates intracellular RNA ligands. Thus, these results further indicate the requirement of RNA ligand binding in Nsp13-mediated cell death and the regulation of this cell death by type I IFNs.

### The SARS-CoV-2 genome shows Z-RNA signatures, binds to Zα domains of ZBP1, and promotes Nsp13-mediated cell death

The role of Nsp13 in promoting cell death, interacting with host RHIM-proteins, and its ability to promote ZBP1 signaling suggest the association of Nsp13 with Z-RNAs. We hypothesize that the SARS-CoV-2 RNA genome consists of base repeats favoring Z-RNA conformation and are generated during its replication process. The repeats of purine and pyrimidine (Pu:Py), predominantly (CG)n, (UG)n/(TG)n, (CA)n, favor Z-RNA/Z-DNA conformation (*30, 72–74*). Also, short interspersed nuclear elements (SINEs) with inverted repeat sequences increase the propensity of RNA to attain Z-conformation (*41, 67, 75, 76*). We have devised a comprehensive analysis, incorporating alternate Pu:Py, inverted tandem repeats, SINEs, and the sequences favoring double-strand RNA formation, for identifying Z-RNA segments in the SARS-CoV-2 genome. We found that Z-RNA-like sequences were distributed across the SARS-CoV-2 genome, but most of these sequences span ORF1a and ORF1b of the genome (**Figure S7A**). We further predicted the RNA secondary structures of these potential Z-RNA hotspots using RNAstructure prediction server (**Figure 7A, S7B**). Among these Z-RNA forming double-stranded RNA (dsRNA) structures, we selected dsRNAs in which alternate Pu:Py repeats stabilized the secondary structure. We identified several SARS-CoV-2 genome hotspots with Z-RNA-favoring sequences using this approach. Recent studies experimentally determined the secondary structure of the SARS-CoV-2 genome in infected cells (*77–79*). The predicted secondary structures of Z-RNA sequences were mapped on the experimentally solved secondary structures of the SARS-CoV-2 genome to check if they retained the same structures as the prediction. Using this secondary structure comparison, the candidate Z-RNAs, which retain the dsRNA conformation in experimentally solved SARS-CoV-2 genome structure, were identified (**Figure 7A, S7B**). The identified Z-RNAs forming secondary structures span the SARS-CoV-2 Nsp1, Nsp2, Nsp3, Nsp4, Nsp6, Nsp13, and Nsp16 coding regions of ORF1ab (**Figure S7B**). By examining the experimentally determined SARS-CoV-2 genome secondary structure, we further identified additional genome sequences with Z-RNA forming propensity in Nsp1 and N-protein coding regions. Together, our rational sequence and structure-based mining of the SARS-CoV-2 genome identified the sequences with high Z-RNA forming propensity.

**Figure 7:**
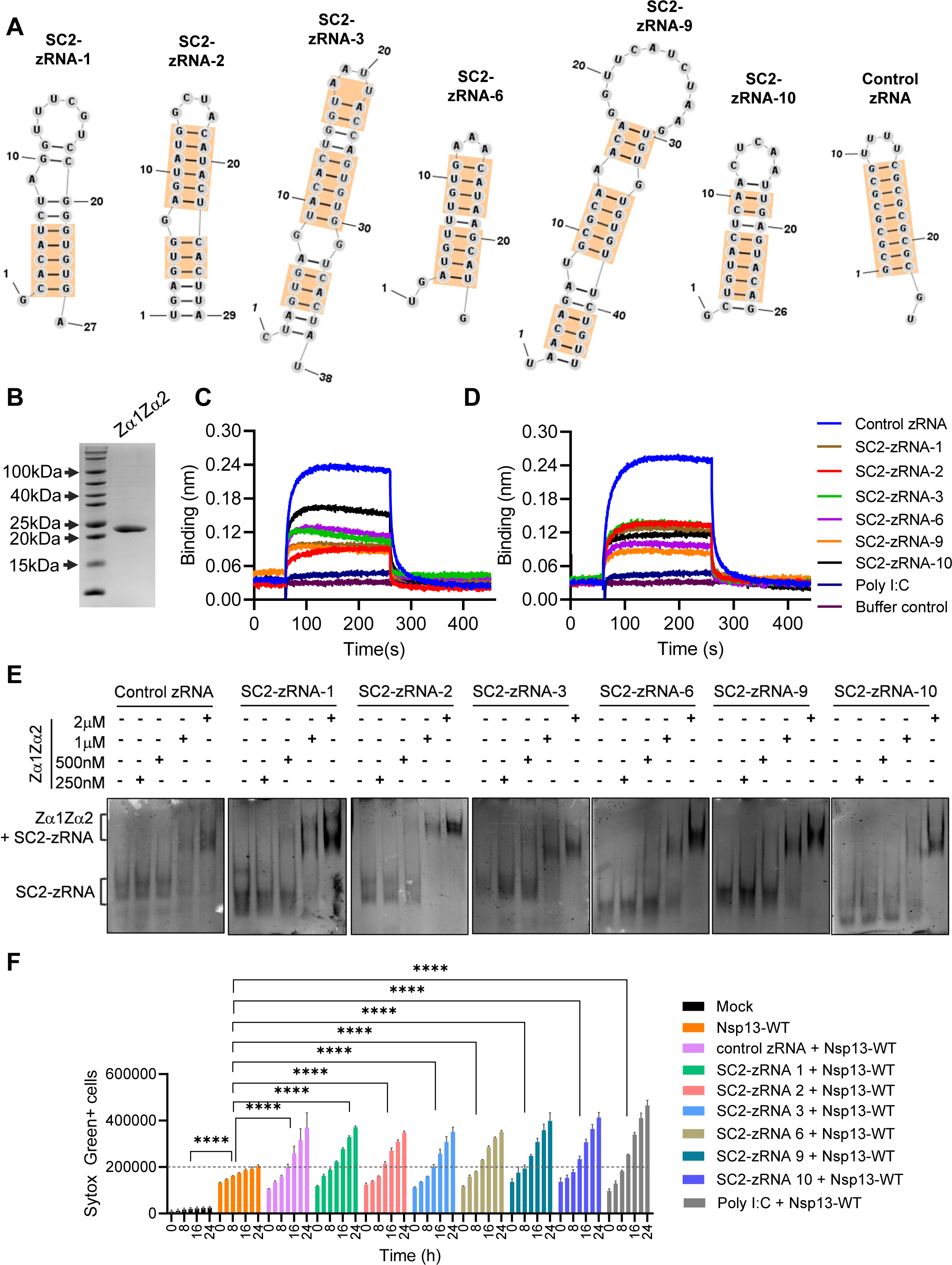
SARS-CoV-2 encode Z-RNA forming genomic segments that enhance Nsp13- mediated cell death. **A,** Specific SARS-CoV-2 genome segments with high Z-RNA forming dsRNA conformations. The orange box outlines represent alternate purine-pyrimidine repeats with Z-RNA forming and Zα-domain binding potential. **B,** SDS-PAGE gel picture representing purified human ZBP1-Zα1α2 domains. **C-D,** Octet binding of SARS-CoV-2 Z-RNAs (SC2-zRNA), control Z-RNA (alternate purine-pyrimidine repeats favoring Z-RNA conformation) and poly(I:C) at 100nM (D) and 200nM (E) concentration to human ZBP1-Zα1α2 domains. **E,** Electrophoretic mobility shift assay of control Z-RNA and SARS-CoV-2 Z-RNAs with human ZBP1-Zα1α2 domains to monitor their interaction. **F,** Real-time cell death measurement by Sytox green staining of Nsp13 expressing L929 cells after transient transfection with control zRNA, SARS-CoV-2 Z-RNAs (SC2-zRNA) or Poly(I:C). \*\*\*\**P*<0.0001 (two way ANOVA, *n*=3). Data shown are mean ± SEM.

To further validate the ability of SARS-CoV-2 Z-RNAs (SC2-zRNA) for binding Z-RNA sensing proteins, we purified Zα domains of human ZBP1 (Zα1-Zα2) to probe its binding to SC2-zRNAs using real-time biolayer interferometry and electrophoretic mobility shift assays (EMSA) (**Figure 7B**). SC2-zRNAs segments were generated by in vitro transcription for binding studies with Zα1-Zα2. We mutated one of the surface-accessible serine residues in Zα2 domain to cysteine (S106C) for labeling purified Zα1-Zα2 protein. Using biolayer interferometry (BLI-Octet) we examined the real-time binding of in vitro transcribed SARS-CoV-2 RNA segments with high Z-RNA propensity (SC2-zRNA-1,2,3,6,9 and 10) to purified Zα1-Zα2 (**Figure 7C,7D**). In this analysis, we included poly(I:C) and a Pu:Py repeat RNA that attain Z-RNA conformation (control zRNA) when incorporated with modified guanine bases (*67, 80*). The control zRNA bound to Zα1-Zα2 protein with high association rates (despite lacking modified guanines) and poly(I:C), and buffer controls bound with very low association rates (**Figure 7C,7D**). Perhaps Zα1-Zα2 binding might have driven the attainment of stable Z-conformation in control zRNA despite lacking modified bases. The SC2-zRNA bound to Zα1-Zα2 protein with higher association rates than poly(I:C) and buffer control but less than the control zRNAs (**Figure 7C,7D**). Notably, the dissociation of control zRNA and SC2-zRNAs from Zα1-Zα2 protein was similar. These observations indicate that the selected SARS-CoV-2 RNA genome segments attain z-RNA conformation and bind to Zα1-Zα2 protein without the requirement of incorporating modified bases in SC2-zRNAs. Furthermore, we performed EMSAs to monitor SC2-zRNAs interaction with Zα1-Zα2 protein. Although we attempted to label Zα1-Zα2 protein by cysteine maleimide labelling for fluorescence imaging of EMSA gels, the labeled protein was degraded rapidly. We further proceeded to use unlabeled protein for EMSA studies. We observed that control zRNA and SC2-zRNAs showed detectable shifts in zRNA sizes, indicating the binding of SC2-zRNAs with Zα1-Zα2 protein (**Figure 7E**). These band shifts were observed at higher Zα1-Zα2 protein concentrations, perhaps because of the faster dissociation rates observed in real-time BEI-Octet analysis (**Figure 7E**). The EMSA experiments further establish that SARS-CoV-2 RNA genome segments attain Z-RNA conformation and bind to Z-RNA sensing Zα domains. Since intracellular RNA ligands regulate Nsp13-mediated cell death, we tested whether SC2-zRNAs promote Nsp13-mediated cell death. Nsp13-expressing L929 cells showed enhanced cell death levels after intracellular delivery of SC2-zRNAs or Poly(I:C) compared to mock-transfected Nsp13-expressing cells (**Figure 7F**). SC2-zRNAs showed a low magnitude of cell death in the absence of Nsp13 expression (**Figure S7C**). These observations suggest that SC2-zRNAs regulate SARS-CoV-2 Nsp13-mediated cell death.

## DISCUSSION

Our findings in this report indicate that Nsp13 and Nsp14 of CoVs harbor putative RHIMs. Viral RHIMs reported to date were found in DNA viruses, and our study characterized viral RHIMs of RNA viruses for the first time. SARS-CoV-2 Nsp13 is a viral RHIM protein promoting human cell death programs. Our analysis further reveals bats as an essential source in the evolution of viral RHIMs in pathogenic RNA viruses. It was intriguing to find that, unlike the viral RHIMs reported to date, SARS-CoV-2 Nsp13 did not inhibit cell death and instead promoted it in specific conditions. A recent study indicated a RHIM-like sequence signature in SARS-CoV-2 Nsp13 based on protein sequence analysis (*81*). Our protein sequence and structure-based analysis of the CoV family revealed CoV-RHIMs in Nsp13 and Nsp14 proteins and their association with bat-originated CoVs. Moreover, the CoV-RHIMs in Nsp13 and Nsp14 showed a distinct pattern of evolution. CoV-RHIM-1 in Nsp13 showed a conserved RHIM sequence pattern in pathogenic human-CoVs and bat-associated beta-CoVs, whereas CoV-RHIMs of Nsp14 are conserved in less pathogenic human-CoVs. Our analysis might facilitate RHIM sequence-based annotation of new bat-CoVs or CoVs that have not infected humans for their possible pathogenic potential. How did Nsp13 and Nsp14 acquire RHIMs mimicking host RHIM-proteins? Bats express RIPK3 and ZBP1 proteins. The conserved CoV-RHIMs in bat-originated CoVs raise the possibility that bat RIPK3 and ZBP1 might have driven the evolution of RHIMs in CoVs to modulate their function. Also, how bats show only mild clinical symptoms despite hosting SARS-CoV-2-like viruses is unknown. RHIM-mediated cell death, the evolution of viral RHIMs, and RHIM-driven virus-host interactions might provide new clues for understanding viral tolerance in bats. Our observations indicate that RNA viral RHIM proteins, like Nsp13, show distinct cell death regulatory mechanisms in bat and human cells. Nsp13 preferentially promotes apoptosis of bat cells, which is less dependent on CoV-RHIM-1 but requires RNA-binding channel function. However, Nsp13 triggers apoptotic and necrotic-like cell death in human cells that depend on CoV-RHIM-1 and RNA binding channel. Thus, bats might have evolved to employ additional mechanisms for modulating viral RHIM-mediated cell death to alleviate viral RHIM-mediated detrimental effects. Our results showing the detectable expression of RHIM proteins in bat cells and their sensitivity to RHIM-dependent cell death triggers suggest that bat cells operate RHIM-mediated signaling albeit less robustly than human cells.

Nsp13 is a helicase protein and a component of the SARS-CoV-2 replication-transcription complex (*43, 54, 55*). SARS-CoV-2 mini replication-transcription complex harbors two Nsp13 molecules in distinct conformation (apo and RNA-bound). The 1B domain of the Nsp13 that constitutes CoV-RHIM-1 is essential for interacting with apo- and RNA-bound Nsp13 molecules in the replication-transcription complex (*54, 55*). The RNA binding channel of Nsp13 comprises domain 1B, Rec1A, and Rec2A and is critical for holding incoming RNA genome. Of note, studies on SARS-CoV-1 and other CoV Nsp13 suggested that alterations or mutations in Nsp13 abolish its helicase activity and restrict viral propagation (*59, 60, 82*). Although the role of Nsp13 in SARS-CoV-2 replication and RNA genome unwinding has been described, its replication-independent functions have not been studied. Our cell death signaling, and biochemical studies indicate a critical role for the CoV-RHIM-1 and RNA-binding channel in Nsp13-mediated cell death. Also, intracellular RNA ligands and SC2-zRNAs further promoted Nsp13-mediated cell death. These observations suggest a replication-transcription complex-independent role of Nsp13 in modulating host responses, and the conformation of the RNA binding channel is critical in this case. Defective viral genomes (DVGs) generated during RNA virus replication are known to affect host innate immune responses (*83, 84*). DVGs are the source of Z-RNAs in IAV infection, and ZBP1 senses DVGs to trigger IAV-induced cell death (*27*). Also, endogenous Z-RNA sensing by ZBP1 in physiological conditions triggers immunopathology (*75, 76, 85, 86*). Our in vitro functional studies revealed a possible ZBP1-like function of Nsp13 to promote programmed cell death. Recent studies show that SARS-CoV-2 infection in vitro and infected patients generate DVGs associated with innate immune activation (*87–89*). Nsp13 might act as a sensor when it is not an integral part of the replication-transcription complex, detect specific RNA ligands formed in infected cells, and trigger cell death like ZBP1. Perhaps, Nsp13 detects specific RNA patterns of the SARS-CoV-2 RNA genome or DVGs that might attain Z-RNA conformation to activate cell death. An intriguing question is why Nsp13 has evolved to promote cell death, unlike other viral RHIM proteins which inhibit cell death? We anticipate that SARS-CoV-2 mediated delayed type I IFN response limits ZBP1-RIPK3 signaling and exerts suboptimal cell death activation at an early stage of infection. Nsp13 might have evolved to trigger RHIM-dependent cell death at later stages of infection to promote virus spread. Nevertheless, future studies need to examine this question. While this manuscript was in preparation, a new study reported that SARS-CoV-2 forms Z-RNAs in infected cells and activates ZBP1-RIPK3 signaling and cell death (*38*). This work demonstrates that ZBP1-RIPK3 signaling promotes lung damage and inflammation during SARS-CoV-2 infection in mice. This study suggests the formation of Z-RNAs in SARS-CoV-2 infection, however, specific RNA genome regions of SARS-CoV-2 attaining Z-RNA conformation and their interaction with Z-nucleic acid binding domains (Zα domains) was unknown. Our stringent pipeline for identifying Z-RNA sequences led us to annotate high-propensity Z-RNA forming sequences in the SARS-CoV-2 genome (SC2-zRNAs). Our biochemical and real-time binding studies demonstrate the binding of SC2-zRNAs to purified Zα1Zα2 domains of human ZBP1 protein. These observations reveal SARS-CoV-2 RNA genome-derived Z-RNA ligands that might activate ZBP1 and Nsp13-mediated cell death.

## METHODS

### Protein sequence and structure analysis

CoV-RHIM sequences in coronaviruses, SARS-CoV-2, and other bat RNA viruses were determined based on well-characterized human and viral RHIM protein sequences and amino acid propensities that favor β-sheet amyloid structures. Coronavirus protein sequence alignments were performed using Clustal Omega multiple sequence alignment tool (from EMBL-EBI). Putative RHIM sequences were compared to annotate variation among coronaviruses. Maximum likelihood phylogenetic analysis of protein sequences was done using PhyML 3.1/3.0aLRT in Phylogene.fr, or MEGA-X and the analysis output was generated in Newick format. Phylogenetic trees were illustrated in MEGA-X (v10.1.7) using PhyML-generated Newick output formats. Crystal structures of Nsp13 (PDB ID: 6ZSL), Apo and RNA-bound forms of Nsp13 (PDB ID: 7CXM), and Nsp14 (PDB ID: 7N0B) of coronaviruses were visualized, and ribbon and surface structure models were prepared using Chimera 1.14 or ChimeraX 1.4 protein model visualization tool.

### Plasmids and constructs

pLVX-EF1alpha-SARS-CoV-2-Nsp13-2xStrep-IRES-Puro,pLVX-EF1alpha-SARS-CoV-Nsp13-2xStrep-IRES-Puro, and pLVX-EF1alpha-MERS-Nsp13-2xStrep-IRES-Puro were a kind gift from Prof. Nevan Krogan’s lab (University of California, San Francisco). Nsp13-Tet-mut (residues 193-196, VQIG→AAAA) and Nsp13-Swap-mut (residues 193-196, VQIG→TVLG) constructs were generated by site-directed mutagenesis using overlap extension PCR. The construct expressing Nsp13-EGFP was generated by subcloning the sequence of EGFP from pLVX-EF1alpha-EGFP- 2xStrep-IRES-Puro to pLVX-EF1a-SARS-CoV-2-Nsp13-2xStrep-IRES-Puro using overlap extension PCR. Nsp13 domain deletion constructs were generated using overlap extension PCRs in the same vector background of the SARS-CoV-2 Nsp13 expression construct. The coding sequence of human ZBP1 (hZBP1) was amplified from pCMV-Human-ZBP1 cDNA clone expression plasmid (Sino Biological Inc., HG19385-UT) and subcloned into pLVX-EF1alpha- 2xStrep-IRES-Puro backbone using EcoRI and BamHI restriction sites. ZBP1-RHIM-Mut (residues 205-208, IQIG→AAAA, N211A) was generated by site-directed mutagenesis using overlap extension PCR. The HA-tag sequence was fused at the C-terminus of ZBP1 through PCR primers. The construct expressing hZBP1-dTomato was generated by fusing dTomato coding sequence at the C-terminus of ZBP1 by overlap extension PCR. Other plasmids used in the study, pcDNA3-HA-RIPK3 (78804) and pcDNA3-FLAG-RIPK1 (78842) were procured from Addgene.

### Cell Culture, transfections, and stimulations

Cell lines used in this study, A549 (human lung carcinoma), HT-29 (human colorectal adenocarcinoma), HCT-116 (human colorectal adenocarcinoma), L929 (mouse connective tissue fibroblast), HEK-293T (human embryonic kidney), Tb1 Lu (bat- *T.brasiliensis* lung epithelial cells) and MDCK (Madin-Darby canine kidney) were obtained from National Centre for Cell Science (NCCS) cell repository and were authenticated by STR profile analysis. Cell lines were tested to be negative for mycoplasma contamination. A549-ACE2 and Vero-E6 cells were obtained from Dr. Shashank Tripathi’s lab (Indian Institute of Science). All the cell lines were cultured at 37°C and 5% CO2 in DMEM growth medium (Thermo Fisher Scientific, 11995040) supplemented with 10% v/v FBS (Thermo Fisher Scientific, 10270106), Antibiotic-Antimycotic (Thermo Fisher Scientific, 15240062) and non-essential amino acids (Thermo Fisher Scientific, 11140-050). Tb1 Lu cells were cultures in EMEM growth medium (Lonza, 12611F), with the conditions mentioned above. For transient expression of proteins, plasmids were transfected into specific cell lines using Xfect (TakaraBio, 631318) or Lipofectamine 2000 (Thermo Fisher Scientific, 11668-019) transfection reagents in reduced serum media Opti-MEM (Thermo Fisher Scientific, 31985070).

For apoptosis and necroptosis activation, HT-29 cells were treated with 30µM Z-VAD(OMe)-FMK (Cayman Chemical, 14463) and 500nM SMAC mimetic SM-164 (MedChemExpress, HY-15989). 2 h after treatment, 100ng ml^-1^ recombinant human TNF-α(Abclonal, RP00001) was added.Tb1 Lu cells were treated with 30µM Z-VAD(OMe)-FMK and 1µM SM-164 2 hours before treatment with 100ng ml^-1^ TNF-α. Necroptosis was inhibited in HT29 and Tb1 Lu cells using 5µM MLKL inhibitor – GW806742X (Sigma Aldrich, SML1990). Influenza A virus (IAV)-infected L929 cells were treated with 30 µM Z-VAD(OMe)-FMK for inducing necroptosis. To activate ZBP1-specific cell death, L929 cells were treated either with a combination of 5ng ml^-1^ Leptomycin-B (Sigma-Aldrich, L2913) and 100ng ml^-1^ IFN-β(Abclonal, RP01076) or with CBL0137 (Cayman Chemicals, 19110). Tb1Lu cells were treated with 5µM CBL0137 alone and in combination with 30 µM Z-VAD.

### IAV infection

Influenza A virus (A/WSN/1933) was generated using 8 plasmid reverse genetics system and propagated in the MDCK cell line to obtain progeny 1 (P1) virus stocks. For IAV infection experiments, cells were seeded in DMEM (Thermo Fisher Scientific, 119905040) supplemented with 10% FBS. 24 h after seeding cells, the media was replaced with DMEM lacking sodium pyruvate (Sigma Aldrich, D6171), and cells were infected with IAV. 2 h post-infection, 10% FBS was added to the cells and real-time cell death analysis was performed using IncuCyte S3 Live-Cell Analysis instrument (Sartorius).

### Lentivirus transduction for generating *Zbp1*-knockout L929 cells and stable transgenic protein expression

All the cell lines were maintained in DMEM containing 10% FBS and 1% Antibiotic-Antimycotic solution. Lentiviral particles expressing Cas9 were generated by transfecting HEK-293T cells with LentiCas9-Blast, psPAX2, and pMD2.G plasmids using Xfect transfection reagent in Opti-MEM media. Lentiviral supernatants were harvested 48 hours post-transfection. L929 cells were then infected with Cas9-lentiviruses in the presence of polybrene (Sigma-Aldrich, TR-1003-G) to obtain cells stably expressing Cas9 protein. The cells were selected with Blasticidin (Thermo Fisher Scientific) and maintained in culture. Lentivirus stocks were prepared using transfer plasmid encoding guide-RNA (gRNA) targeting Zbp1 (Sequence: GTCCTTTACCGCCTGAAGA). L929 cells stably expressing Cas9 were infected with Zbp1 gRNA-lentiviruses in the presence of polybrene and selected with Puromycin (Sigma-Aldrich). Loss of ZBP1 protein expression after the selection was confirmed by immunoblotting analysis with anti-ZBP1 antibody.

To stably express Nsp13 and other domain deletion constructs of Nsp13 in A549, HCT-116, HT-29 and Tb1 Lu cells, lentivirus particles for transducing the cells were generated in HEK293T cells by transfecting Nsp13 expression constructs along with pCMV-VSVG and psPAX2 plasmids. Lentivirus supernatants were harvested 48 h after transfection, supplemented with an additional 10% FBS and stored at −80°C. The cells were transduced with lentivirus in the presence of 12mg ml^-1^ of polybrene and subjected to selection with Puromycin (Sigma-Aldrich, P8833).

### SARS-CoV-2 Infection and virus titer assays

Constructs expressing SARS-CoV-2 Nsp13-WT, Nsp13-Tet-Mut, Δ1B-Rec1A-Rec2a, Δ1B were transfected in to A549 cells that express ACE2 protein (A549-ACE2). 24 h after transfection, cells were infected with the SARS-CoV-2 virus (Isolate Hong Kong/VM20001061/2020; NR-52282; BEI Resources, NIAID, NIH) at an MOI of 3. Virus inoculum was prepared in DMEM (Thermo Fisher Scientific, 119905040) supplemented with 2% FBS and allowed for 1 hour at 37°C for virus adsorption onto cells. After 1 h, DMEM containing 2% FBS was added to cells. 48 h after infection, cells were fixed using 4% formaldehyde and subjected to SYTOX green staining for cell death analysis. Whole cell lysates were collected for immunoblotting analysis of SARS-CoV-2 N-protein.

Viral titers were estimated by Vero E6 cell-based plaque assay. At 90% confluency after seeding, cells were incubated at 37°C, with 150 μl of increasing dilutions of virus supernatant collected from SARS-CoV-2 infected A549-ACE2 cells transiently expressing Nsp13-WT, Nsp13-Tet-Mut, Δ1B-Rec1A-Rec2A, Δ1B proteins. The virus inoculum was removed after 1 h and the cells were overlaid with 0.6% Avicel (Dupont, RC-591) prepared in DMEM containing 2% HI-FBS. After 48 h of incubation at 37°C, the cells were fixed with 4% paraformaldehyde. Following fixation, the cells were stained with Crystal Violet (Sigma Aldrich, C0775) for 20 minutes, and plaques were visualized.

### Ethics statement

This study was performed according to the guidelines and in compliance with institutional biosafety guidelines (IBSC/IISc/KS/11/2020; IBSC/IISc/KS/32/2021). SARS-CoV-2 experiments were conducted at viral Biosafety level-3 facility and IAV-WSN experiments were conducted at Biosafety level-2 facility at Indian Institute of Science (IISc).

### Real time cell death analysis

Real-time cell-death assays were performed using a two-color IncuCyte S3 Live-Cell Analysis instrument (Sartorius). Cell lines were seeded in 12 well plates and treated with cell death stimulating agents or infected with viruses. Dead cells were stained with 20nM Sytox Green (Thermo Fisher Scientific, S7020), a cell-impermeable DNA-binding fluorescent dye which rapidly enters dying cells after membrane permeabilization and fluoresces green. For quantifying cells undergoing apoptosis, 2.5 µM Caspase-3/7 Green Dye (Sartorius, 4440) was used. The incucyte Caspase-3/7 dye is a DNA intercalating dye that is activated on cleavage by Caspase3/7. The resulting images and the fluorescence signals were analysed using IncuCyte S3 software, which provides a count for the Sytox or Cleaved Caspase-3/7 Green-positive cells using microscopic images. This data was plotted using GraphPad Prism 9.0 software.

### Immunoprecipitation studies for probing Nsp13 association with host RHIM-proteins

HEK-293T cells were seeded in 100mm dishes and transfected with the constructs expressing WT and mutant forms of Nsp13, ZBP1and RIPK3 for co-immunoprecipitation studies. Depending on the cell death, 27-48 hrs after transfection the cells were lysed in NP-40 lysis buffer (1.0 % NP40, 50mM Tris, pH 8.0, 150mM NaCl supplemented with protease inhibitor and phosphatase inhibitor cocktails), and lysates were cleared by centrifugation at 12000 rpm for 20 min. The cell lysates were incubated with 10 μg of anti-Streptag-Antibody and 2 μg of the indicated primary antibodies for 4–5 h at 4°C. Protein A/G PLUS-Agarose (Santa Cruz Biotechnology, 38221990) beads were washed with 1XPBS, incubated for 30 min with 3% BSA for blocking, and added to the samples. After 2 h of incubation at 4°C, the agarose beads were then collected by centrifugation at 1000 rpm and washed three times with 1:1 PBS and NP-40 lysis buffer. Immunoprecipitates were eluted from the beads in the sample buffer and subjected to immunoblotting analysis.

The separated insoluble (pellet) fractions were dissolved in NP-40 lysis buffer containing 2% SDS. After solubilization and centrifugation, the supernatant was incubated with antibodies and samples were processed as mentioned for soluble fractions (above).

### Capturing oligomeric complexes in soluble and insoluble fractions of cellular lysates

HEK-293T cells transiently transfected with indicated clones were subjected to lysis with NP-40 lysis buffer (1.0% NP40, 50mM HEPES, pH 8.0, 150mM NaCl with protease inhibitor and phosphatase inhibitor cocktails) containing 2mM Pierce Premium Grade DSP (Thermo Fisher Scientific, PG82081). Whole cell lysates were collected and incubated at room temperature for 45 min. The reaction was stopped by adding 1M Tris-HCl and incubating for 15 minutes. Whole cell lysates were centrifuged at 10,000 rpm for 15–20 min to separate insoluble fractions (pellet) and soluble fractions. Insoluble fractions (pellets) were washed using lysis buffer and dissolved in NP-40 lysis buffer. Soluble and insoluble lysates were mixed with 4x sample buffer with or without β-mercaptoethanol (BME) and subjected to SDS-PAGE and immunoblotting analysis to monitor the oligomeric status of complexes.

### Immunoblotting analysis

The cells were lysed using NP40-lysis buffer and added sample-loading buffer containing SDS and BME. The lysates were resolved in 8-12% SDS-PAGE gels, and the gels were subjected to transfer onto the PVDF membrane. The membranes were blocked in 5% skimmed milk at room temperature for 1 h and further incubated with primary antibodies overnight at 4°C. The membranes were washed using Tris-buffered saline with tween 20 (TBST), subjected to horseradish peroxidase (HRP)-conjugated secondary antibodies for 1h at room temperature, and then washed with TBST. The primary antibodies used in the study were anti-ZBP1 (Adipogen, AG-20B-0010-C100), anti-ZBP1 (Santa Cruz, sc-271483), anti-Strep Tag (Qiagen 34850), anti-MLKL (Cell Signalling Technology, 37705, Mouse-specific), anti-MLKL (Cell Signalling Technology, 14993, Human-specific), anti-p-MLKL (Cell Signaling Technology, 37333, Mouse-specific), anti-p-MLKL (Cell Signaling Technology, 91689, Human-specific), anti-RIPK3 (Cell Signaling Technology, 95702, Mouse-specific), anti-RIPK3 (Cell Signaling Technology, 10188, Human-specific) anti-IAV N-Protein (Invitrogen, PA5-32242), anti-CASP3 (Cell Signaling Technology, 9662), anti-HA Tag (Invitrogen,26183), anti-Flag (Invitrogen, MA1-91878),anti-RIPK1 (Cell Signaling Technology, 3493), anti-GAPDH (Invitrogen,15738), anti-SARS-CoV-2 N protein (Cell Signaling Technology, 33717). HRP-conjugated secondary antibodies used in this study were Jackson Immuno Research Laboratories anti-rabbit (111-035-047) or anti-mouse (315-035-047) antibodies. Blots were developed using Immobilon Forte Western HRP substrate (Millipore, WBLUF0500) and visualized using Image Quant LAS500 or Image Quant 800 (Cytiva, Amersham).

### Immunofluorescence and DNA-PAINT imaging

HEK-293T cells were transfected with the respective plasmids (Zbp1-HA, ZBP1-dTomato, ZBP1-RHIM-Mut-HA, RIPK1-FLAG, and RIPK3-HA) along with Nsp13-EGFP. For checking RHIM-dependent interaction, HEK-293T were further transfected with Nsp13-WT and Nsp13-Tet-Mut in combination with RIPK3-HA. 20-22 h after transfection, the cells were fixed with 4% paraformaldehyde in 1X PBS (ChemCruz, sc-281692) for 20 minutes at room temperature. After fixing, cells were permeabilized using 0.1% Triton X-100 for 10 min at room temperature. Nonspecific binding was blocked using 3% BSA (GBiosciences, RC1021). Cells were incubated with anti-HA Tag (Invitrogen, 26183) or anti-FLAG Tag antibody for 1 h at room temperature. The cells were then subjected to staining with Alexa Fluor 568-conjugated anti-mouse IgG (Invitrogen, A-11004; 1:500) for 30 mins at room temperature. After each step, the cells were washed twice with 1X PBS. The cells were counterstained with DAPI. All images were acquired using Olympus FV 300 Confocal microscope system. The colocalization analysis was performed using ‘Coloc2’ plugin of Fiji. The plugin was used to quantify Pearson’s correlation coefficient as a mathematical measure of colocalization.For DNA points accumulation for imaging in nanoscale topography (DNA-PAINT) imaging, HEK-293T cells were seeded and co-transfected with constructs expressing Nsp13-EGFP along with ZBP1-HA and RIPK3-HA separately. 20-22 h after transfection, the cells were fixed for 15 min using 4% paraformaldehyde (Electron Microscopy Sciences, 15710), preheated to 37°C and subjected to 5 washes with PBS, pH 7.4. 1 mg ml^-1^ sodium borohydride solution in PBS was used to quench the free aldehyde groups, followed by 5 washes with PBS. The cells were then permeabilized with 0.5% Triton X-100 in PBS for 15 min, followed by 2 washes with PBS. The cells were then incubated with 3 % Bovine Serum Albumin (Sigma, A4503) in PBS for 45 min for blocking, followed by 2 washes with PBS. 1:200 dilution of anti-GFP (Invitrogen, A-11122) and anti-HA tag (Invitrogen, 26183) primary antibodies were prepared in 3 % Bovine Serum Albumin, added to the wells, and incubated overnight in the dark at 4°C followed by 2 washes with PBS. 1:200 dilution of the docking-strand conjugated anti-mouse and anti-rabbit secondary antibody in 3% Bovine Serum Albumin was added to the wells and incubated for 45 minutes at room temperature, followed by 2 washes with PBS. Gold nanoparticles diluted at a 1:5 ratio in PBS was added to the chamber and incubated for 5 min followed by 5 washes with PBS.

Microscopy was performed on a Nikon Ti-2 eclipse microscope equipped with a motorized H-TIRF, perfect focus system (PFS), and a Teledyne Photometrics PRIME BSI sCMOS camera. Illumination using 561 nm wavelength lasers was done using the L6cc Laser combiner from Oxxius Inc., France. Imaging was done under Total Internal Reflection conditions. Imager sequences used for DNA-PAINT were 5’-AGGAGGA-Cy3B-3’ and 5’-AGAGAGA-Cy3B-3’. For 2-color imaging of Nsp13-EGFP and RIPK3-HA or ZBP1-HA, the imager solution from the first round of imaging was washed 5 times with buffer C (1× PBS and 500 mM NaCl) with 2 min of incubation. The second imager solution was prepared in buffer C, added to the wells, and imaged at the same plane. For all DNA-PAINT imaging rounds, the laser power was set at 313W/cm^2^ on the imaging plane, and the image acquisition rate was set to 10 Hz in HiLo mode. The obtained raw fluorescence data was reconstructed using ‘Localize’ tool embedded in the Picasso Software suite to obtain a super-resolved image (*90*). Drift correction was performed by redundant cross-correlation (RCC) followed by correction with fiducial markers. RCC was also used to align the super-resolved structures from the two imaging channels. Reconstructed data was rendered in Picasso:Render. For 2-color data, reconstructed files for both rounds of imaging were individually drift-corrected and loaded into ‘Render’ together using RCC or gold nanoparticles to align the two images.

### Protein Purification

The bacterial codon-optimized coding sequence of ZBP1-Zα1Zα2 domains was synthesized (Thermo Fisher Scientific) and subcloned into N terminally His-Tagged pNIC-ZB vector. The plasmid expressing pNIC-ZB-ZBP1-Zα1Zα2 was transformed into Rosetta DE3 (kind gift from Prof. B Gopal from MBU, IISc) and grown overnight on LB agar plates with kanamycin and chloramphenicol. Single colonies from the plate were inoculated 20ml LB media and incubated overnight at 37°C at 180 rpm. 20ml overnight cultures were inoculated into 1000 ml of LB containing kanamycin and chloramphenicol. Cultures were grown at 37°C, induced at 0.6 OD (600nm) by adding 0.3 mM IPTG, then incubated for 4 h at 37°C. The pellet lysed in the lysis buffer 50 mM HEPES (pH 7.5), 500 mM NaCl, 5 % glycerol, 1 mM PMSF,1 mM DTT and 5 mM imidazole). Cells were disrupted by subsequent sonication until a clear solution was obtained. The cells were further centrifuged at 18300g for 40 min at 4°C. Ni-NTA beads (Gbioscieces, 786-940) were loaded on a gravity flow column and washed with 50 ml of lysis buffer containing 10 mM imidazole and a further wash with 25 ml of buffer containing 1M NaCl. Proteins were then eluted with gradient elution (10 mM Tris HCl, 500 mM NaCl, 5% glycerol, 1 mM PMSF, and no imidazole to 500 mM imidazole). The eluted protein was run on a 15% SDS-PAGE gel to check the purity. The eluted protein fraction was further treated with TEV protease to exclude the Z-basic part from the purified protein. Dialysis was performed for the eluted product with 20 mM Tris 7.5, 50 mM NaCl, 2 mM DTT overnight at 4°C. The eluted fraction was passed through the Heparin column. The column was washed with 50 ml of 50 mM HEPES (pH 7.5), 50 mM NaCl, 5% glycerol, 1 mM PMSF. The eluted fractions were pooled and diluted in no salt buffer (50 mM HEPES (pH 7.5), 5 % glycerol,1 mM PMSF). These fractions were further loaded onto the Resource-S cation exchange column (Cytiva, 17117801), which was equilibrated with start buffer (20 mM HEPES pH 7.5 and 50 mM NaCl). The bound protein was eluted using gradient elution buffer (50 mM to 1 M NaCl). The eluted fractions were run on SDS-PAGE to check the purity of the protein (MW ∼22.8 kDa), and the fractions that have pure proteins were pooled, snap frozen, and stored at −80°C.

### SARS-CoV-2 Z-RNA predictions

The SARS-CoV-2 Wuhan-Hu-1 RNA genome (NCBI, NC_045512.2) was used for predicting the possible Z-RNA forming sequences. Alternative purine-pyrimidine (Pu:Py) repeats and short interspersed nuclear elements (SINEs) with inverted repeats favor RNAs to attain Z-conformation (*27, 41, 67, 73, 80, 91, 92*). SARS-CoV-2 genome was assessed for the presence of Pu:Py repeats and SINE-like sequences using Microsatellite repeat finder, the non-B DNA Motif Search Tool (nBMST) and RNAfold developed by Mathew’s lab for predicting the secondary structures of RNAs (*93, 94*). In the Microsatellite repeat finder, the parameters were extended to inverted repeats, tandem repeats, and Z-DNA-like sequences for identifying Z-RNA-like features. The repeat sequence length was set to a maximum 6 bases, as previous studies established that the Z-nucleic acid sequences are repeats of Pu:Py with a maximum length of 6 (*91, 92, 95, 96*). Further, the alternative Pu:Py repeats spanning inverted repeats in the SARS-CoV-2 genome were selected, and the secondary structure was predicted using RNAfold. The nBMST was executed to delineate the sequences forming alternative conformations that differ from the canonical right-handed Watson-Crick double-helix. By utilizing the tool, we predicted sequences with inverted short tandem repeats that form slipped/hairpin-like structures and Z-DNA-like motif sequences with repeats of purines and pyrimidines. The secondary structure of all the possible inverted repeats in the SARS-CoV-2 genome identified as mentioned above was predicted using RNAfold server to illustrate the canonical base pairs that RNA can attain considering free energy minima. The secondary structure of the SARS-CoV-2 genome in infected cells has been solved recently (*77–79*). Using this experimentally solved complete SARS-CoV-2 genome as a reference, we compared the secondary structures of predicted Z-RNA favoring inverted repeats with the experimentally solved secondary structure and precise base pairing events of the SARS-CoV-2 genome. Following the annotations, the SARS-CoV-2 RNA genome segments with Z-RNA forming potential were selected that retain secondary structures and base pairing like experimentally solved structures.

### EMSA for probing ZBP1-Zα1Zα2 association with SARS-CoV-2 z-RNA fragments

To perform EMSAs, SARS-CoV-2 Z-RNA (SC2-zRNA) fragments were generated using in vitro transcription method. The HiScribe T7 High Yield RNA Synthesis Kit (NEB, E2040S) was used to transcribe the template DNA per the manufacturer’s instructions. The SARS-CoV-2 RNA fragments with Z-RNA forming potential were synthesized as DNA oligos with an upstream T7 promoter sequence (GGATCCTAATACGACTCACTATA). To synthesize uncapped RNA, 1 μg of DNA containing the T7 promoter upstream was mixed with the provided reaction buffer and the DNA oligos complementary to the T7 promoter. The mixture was subjected to a thermal cycling step at 95°C for 5 min. Following the thermal cycling, NTPs (provided in the kit) for RNA synthesis were added to the reaction mixture. The specific NTPs and their concentrations are typically included in the HiScribe T7 High Yield RNA Synthesis Kit. The reaction mixture containing the DNA template, reaction buffer, NTPs, and T7 RNA polymerase was then incubated at 37°C for 14-16 h. This extended incubation period allows the T7 RNA polymerase from the kit to catalyze the transcription process, resulting in the synthesis of uncapped RNA. The purified ZBP1-Zα1Zα2 domain was incubated at different concentrations with different SC2-zRNAs for 30 min at 23°C in the RNA binding buffer (10 mM HEPES pH 7.5, 50 mM NaCl). The RNA-protein mixture was then loaded onto the 6% Native PAGE gel and was run in Tris-boric acid-EDTA (TBE) buffer at 120 V for 40 min at 4°C. Once electrophoresis was completed, the gels were post-stained with ethidium bromide and imaged in the Bio-Print VILBER imaging system.

### SC2-zRNA transfections

24 h after seeding, L929 cells were transfected with the construct expressing Nsp13-WT. 20 h after transfection, the cells were transfected with specified RNAs. 5 µg of RNA (poly I:C, Positive control Z-RNA, indicated SARS-CoV-2 Z-RNAs) was incubated with Lipofectamine 2000 transfection reagent (Invitrogen, 11668019), for 15 mins at room temperature. After incubation, the transfection mix was added to the cells in reduced serum media Opti-MEM, and cells were subjected to real-time cell death analysis using Sytox Green staining in Incucyte.

### Biolayer interferometry (BLI)-Octet

The binding kinetics of ZBP1-Zα1Zα2 with SARS-CoV-2 RNAs were evaluated using the Octet RED96 instrument (Sartorius). First, amine-reactive second-generation (AR2G) biosensors were equilibrated in a buffer PBS pH 7.4. Then, the AR2G biosensor channel was activated using EDC and sulfo-NHS reagents (Sigma). Following this, ZBP1-Zα1Zα2 was diluted to a concentration of 10 µg ml^-1^ and immobilized on activated AR2G biosensors for 300 sec in a 10 mM sodium acetate buffer, pH 4.0. The reaction was quenched by utilizing an excess of sulfo-NHS esters in 1 M ethanolamine. For monitoring SC2-zRNAs interaction with ZBP1-Zα1Zα2, black 96-well plates (Nunc F96 Microwell, Thermo Fisher Scientific) were loaded with 200 μl of the protein or the buffer, maintained at 25°C, and agitated at 1000 rpm. The baseline for each read was recorded using the buffer alone. The association of ZBP1-Zα1Zα2 with SC2-zRNAs at 100 nM and 200 nM concentration was captured for 300 sec, followed by the dissociation for 200 sec by dipping the sensor into the buffer without protein. After each kinetic assay, the biosensor chip was regenerated using 0.1 M Glycine-HCl. The acquired kinetic data was analyzed using the manufacturer’s software (Data Analysis HT v11.1). A global fitting approach was employed to analyze the data, to fit specific sensograms with a 1:1 binding model.

## ACKNOWLEDGEMENTS

We thank S.K. lab members for their critical comments. We are very grateful to Nevan Krogan, Margaret Soucheray, and Ujjwal Rathore from the University of California San Francisco for providing SARS-CoV-2, SARS-CoV and MERS-CoV protein expression clones. We thank Paul G. Thomas from St. Jude Children’s Research Hospital for the helpful discussions. We thank Raghavan Varadarajan and Simran Srivastava from Molecular Biophysics Unit, IISc, for helping with Octet-BLI experiments. This work was supported by funding from the Science and Engineering Research Board (SERB-DST) (SRG/2021/000632) (EEQ/2021/000274), Department of Biotechnology (DBT) (BT/PR45145/COT/142/24/2022), Indian Council of Medical Research (ICMR) (2021-14148/CMB/ADHOC-BMS), the Indian Institute of Science (IISc) Start-up grant, DST-FIST infrastructure fund and Infosys Foundation, Bengaluru, India.

## AUTHOR CONTRIBUTIONS

S.K. conceptualized the study; S.M., D.J., A.A.D., S.N., and S.K. designed experiments, performed cell culture and biochemical experiments; S.K., K.B., and A.A.D. performed sequence and phylogenetic analysis; D.J. and S.K. performed Z-RNA mining and sequence analysis; D.J. and M.S. performed in vitro transcriptions, purification of Za protein, EMSA, and Octet binding studies; S.M, A.A.D., and M.A. performed imaging and DNA-PAINT studies; S.M., O.K., and A.A.D. performed SARS-CoV-2 infections and viral titer assays; S.T. assisted with SARS-CoV-2 infection experiments; M.G. helped with protein purification and DNA-PAINT experiments; S.M., D.J., A.A.D, and S.K. wrote the manuscript; M.G. and S.K. supervised the study; S.K. provided the guidance, and brought the funding; All the authors contributed in manuscript editing.

## SUPPLEMENTARY FIGURES

**Figure S1: Structure-guided domain deletions in Nsp13 suggested the role of RNA binding channel and CoV-RHIM in promoting cell death**. **A,** The secondary structures and surface accessibility (ribbon and surface structures) of CoV-RHIMs in SARS-CoV-2 Nsp13 and Nsp14. SARS-CoV-2 Nsp13 (PDB ID: 6ZSL) (dark gray) and the viral RHIM-1 (green); SARS-CoV-2 Nsp14 (PDB ID: 7N0B, chain-B) (blue), viral RHIM-2 (green), and viral RHIM-3 (orange). **B,** Schematic of the Nsp13 and domain deletion constructs generated for studying Nsp13 mediated cell death. **C,** Immunoblot analysis of lysates of HEK-293T cells indicating the protein expression of Nsp13 domain deletion constructs. **D,** Microscopic analysis of HCT-116 and A549 cells expressing Nsp13-WT and domain deletion constructs after lentivirus infection and puromycin treatment. Scale bar, 50μm. **E & F,** Real-time cell death measurement by Caspase-3/7 and Sytox green staining of HCT-116 cells infected with lentiviruses expressing SARS-CoV-2 Nsp13-WT, Nsp13-Tet-mut, Nsp13-Δ1B and Nsp13-Δ1B-Rec1A-Rec2A constructs. **G**, Real-time cell death measurement by Sytox green staining of HCT-116 cells ectopically expressing SARS-CoV-2 Nsp13 constructs as indicated. \*\*\*\**P*<0.0001, \*\**P*=0.0046, \**P*=0.0178 (two way ANOVA for the 36 hr time point, *n*=3). Data shown are mean ± SEM. **H,** Immunoblot analysis of SARS-CoV-2 nucleocapsid (N) protein and GAPDH in A549-ACE2 cells expressing Nsp13 and its mutant constructs infected with SARS-CoV-2.

**Figure S2: Structure prediction and comparison of Nsp13-WT and Nsp13-Tet-mut indicated local conformational differences: A,** Nsp13 protomers in apo or RNA bound conformation in SARS-CoV-2 mini replication-transcription complex structure (PDB ID: 7CXM). **B-D,** Structure analysis of Nsp13 Apo, RNA bound and Tet-mut conformations to indicate domain-specific conformation perturbations. **B,** Structure alignment of Apo-Nsp13 and Nsp13-RNA bound conformation. **C,** Structure alignment of Apo-Nsp13 and Nsp13-Tet-mut. **D,** Structure alignment of Nsp13-RNA bound and Nsp13-Tet-mut. **E-G,** Alanine mutations in core tetrad residues of Nsp13 alter RNA binding channel conformation and abolish Nsp13-mediated cell death. **E,** The structure of RNA bound Nsp13 showing open RNA binding channel conformation (grey colored ellipse and yellow surface structure). **F,** The structure of Apo-Nsp13 showing closed RNA binding channel conformation. Apo and RNA-bound Nsp13 structures were aligned to show the displacement of 1B, Rec1a and Rec2a domains at the RNA binding channel. **G,** The predicted structure of Nsp13-Tet-mut shows altered RNA binding channel conformation. Alignment of Nsp13-Tet-mut and Nsp13-RNA bound structures showing the open RNA binding channel conformation of Nsp13-Tet-mut.

**Figure S3: Nsp13 promotes bat cell death: A,** Real-time cell death measurement by Sytox green staining of Tb1 Lu cells transiently transfected with 500ng, 1μg or 1.5μg of Nsp13-WT. **B,** Immunoblot analysis of lysates from Tb1 Lu cells showing expression of Nsp13 WT, Nsp13-Tet-Mut, Δ1B-Rec1A-Rec2A or Δ1B after transient transfection. **C,** Representative cell death images of Tb1 Lu cells stained with Sytox after transfection with the constructs as in panel-B, acquired using Incucyte imaging analysis system**. D,** Real-time cell death measurement by Sytox green staining of Tb1 Lu cells expressing the constructs as in panel-B**. E & F,** Real-time cell death measurement by Sytox green staining of HT-29 (**E**) and Tb1 Lu cells (**F**), after treatment of TNF, zVAD and SMACmimetic (SMACmim) in the presence of MLKL inhibitor (MLKL-Inh). \*\*\*\**P*<0.0001 (two way ANOVA, n=3), ns – not significant two way ANOVA). **G,** Immunoblot analysis of lysates from Tb1 Lu cells treated with TNF, SMACmim, z-VAD alone or in indicated combinations, showing expression of MLKL, pMLKL and GAPDH.

**Figure S4: Nsp13 associates with host-RHIM proteins through RHIM-mediated interactions. A,** Real-time cell death measurement by Sytox green staining of HT-29 cells transiently transfected with 500ng or 1μg of Nsp13-WT, Nsp13-Tet-mut and Nsp13-Swap-mut constructs. TNF and SMACmimetic (SMACmim) was used to induce apoptosis, and TNF, zVAD and SMACmim was used to induce necroptosis. **B,** Schematic of Nsp13 tagged with EGFP at its C-terminus. **C,** Immunoblot analysis of HEK-293T cell lysates indicating the expression of Nsp13-WT and Nsp13-EGFP. **D,** Microscopic analysis of A549 and HCT-116 cells infected with lentiviruses expressing SARS-CoV-2 Nsp13-WT, Nsp13-EGFP or EGFP followed by puromycin treatment. Scale bar, 50 μm. **E,** Confocal microscopy imaging of HEK-293T cells expressing RIPK1-FLAG or Nsp13-EGFP and RIPK1-FLAG. Scale bar, 10μm. **F,** Representative Incucyte images showing the real-time colocalization of Nsp13-EGFP and ZBP1-dTom upon expression in HEK-293T cells. Scale bar, 60 μm. **G,** Confocal microscopy imaging of HEK-293T cells expressing ZBP1-WT, ZBP1-RHIM-mut or Nsp13-EGFP alone and Nsp13-EGFP in combination with ZBP1-WT or ZBP1-RHIM-mut. Scale bar, 10μm. **H,** Quantitation of colocalization signals using Pearson’s coefficient of colocalization for the microscopic images of HEK-293 cells expressing Nsp13-EGFP in combination with ZBP1-WT or ZBP1-RHIM-mut. **I,** Immunoblot analysis of anti-Strep-Tag (Nsp13) and anti-HA-tag (RIPK3) immunoprecipitates from soluble fractions of HEK-293T cell lysates expressing Nsp13-WT, Nsp13-Tet-Mut and RIPK3 individually or in indicated combinations of Nsp13 and RIPK3 constructs.

**Figure S5: Nsp13 assembles into large complexes and modulates host RHIM protein complexes and cell death activation**. **A,** Immunoblot analysis of crosslinked lysates of HEK-293T cells (soluble and insoluble fractions) expressing SARS-CoV-2 Nsp13-WT, Nsp13-Tet-mut, Nsp13-Swap-mut, ZBP1 or co-expressing Nsp13-WT + ZBP1 and Nsp13-Tet-mut + ZBP1. O-oligomer complexes; M-Monomer. **B & C,** DNA-PAINT images of SARS-CoV-2 Nsp13 and ZBP1 or RIPK3 in HEK-293T cells. Scale bars, 200nm. **D,** Immunoblot analysis of phosphorylated MLKL (p-MLKL), ZBP1, IAV nucleoprotein (IAV-NP) and pro and cleaved form of Caspase-3 (CASP3) in WT and *Zbp1*^-/-^ L929 cells 24 h after mock or IAV or IAV+zVAD infection.

**Figure S6: Nsp13-mediated cell death is modulated by IFN-β and the intracellular RNA ligands**. **A,** Representative cell death images acquired by Incucyte imaging analysis system treated with IFN-β and poly(I:C) in mock or Nsp13 transfected L929 cells. Scale bar, 200μm. **B-D**, Real-time analysis of cell death by Sytox green staining of mock or Nsp13 transfected L929 cells treated with IFN-β (**B**), intracellular delivered PolyI:C (transfection) (**C**) and ectopic treatment (tr) or intracellular delivered Poly(I:C) (trn-transfection) (**D**).

**Figure S7: Predicted SARS-CoV-2 Z-RNAs and their role in promoting Nsp13-mediated cell death**. **A,** Schematic representation of SARS-CoV-2 genome organization and hot spots of the genome with Z-RNA conformation forming potential. **B,** The list of predicted Z-RNA sequences and their location in the SARS-CoV-2 RNA genome. **C,** Real-time cell death measurement by Sytox green staining of L929 cells after transient transfection with control zRNA, SARS-CoV-2 Z-RNAs (SC2-zRNA) or Poly(I:C) without Nsp13.

