## Supplementary figures for "Bat RNA viruses employ viral RHIMs orchestrating species-specific cell death programs linked to Z-RNA sensing and ZBP1-RIPK3 signaling"

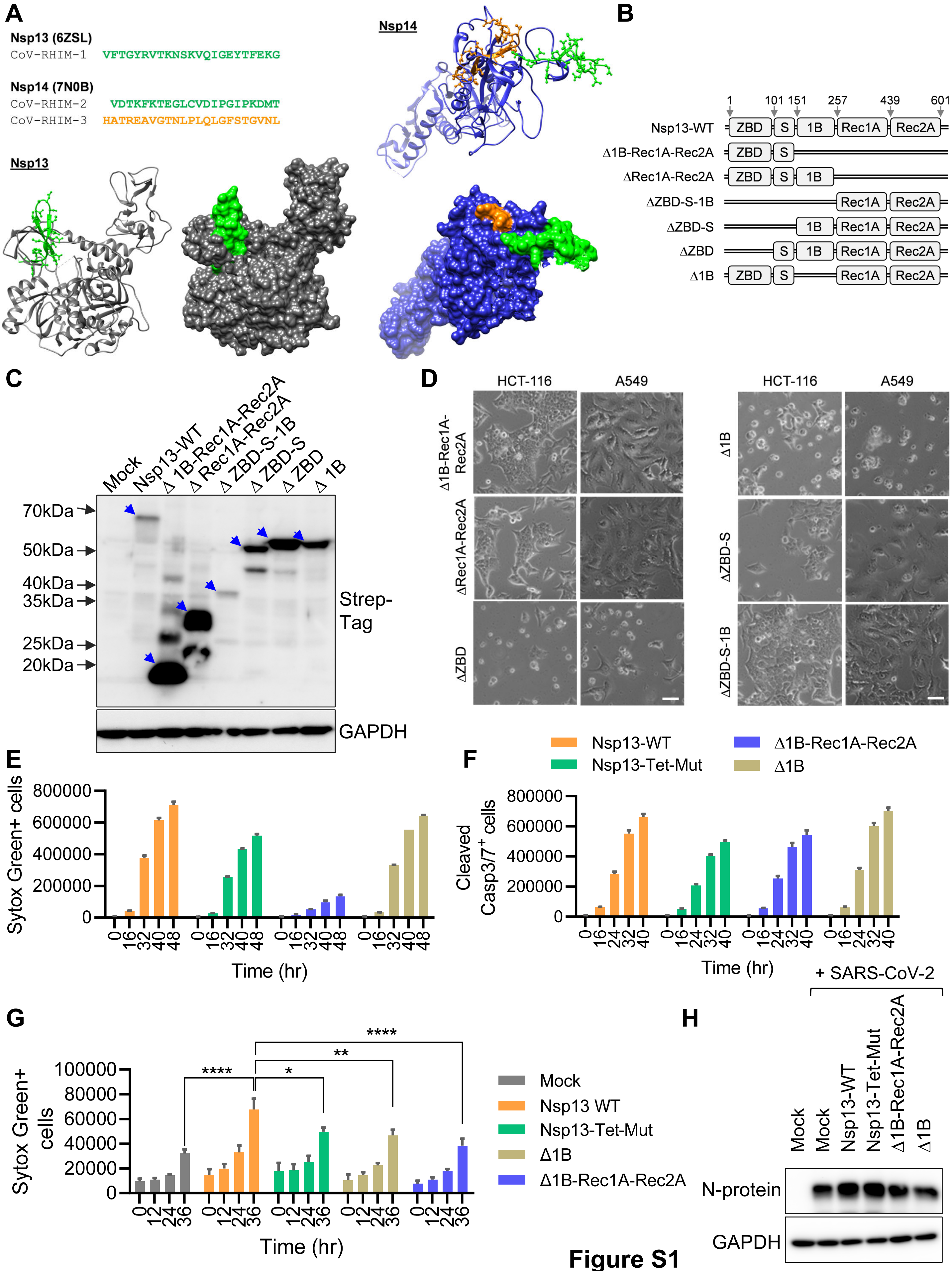
A
Nsp13 (6ZSL)
CoV-RHIM-1 VFTGYRVTKNSKVQIGEYTFEKG
Nsp14 (7N0B)
CoV-RHIM-2 VDTKFKTEGLCVDIPGIPKDMT
CoV-RHIM-3 HATREAVGTNLPLQLGFSTGVNL
Nsp13
Nsp14
B
1 101 151 257 439 601
Nsp13-WT ZBD S 1B Rec1A Rec2A
Δ1B-Rec1A-Rec2A ZBD S
ΔRec1A-Rec2A ZBD S 1B
ΔZBD-S-1B Rec1A Rec2A
ΔZBD-S 1B Rec1A Rec2A
ΔZBD S 1B Rec1A Rec2A
Δ1B ZBD S Rec1A Rec2A
C
Mock
Nsp13-WT
Δ 1B-Rec1A-Rec2A
Δ Rec1A-Rec2A
Δ ZBD-S-1B
Δ ZBD-S
Δ ZBD
Δ 1B
70kDa
50kDa
40kDa
35kDa
25kDa
20kDa
Strep-Tag
GAPDH
D
HCT-116
A549
Δ1B-Rec1A-Rec2A
ΔRec1A-Rec2A
ΔZBD
Δ1B
ΔZBD-S
ΔZBD-S-1B
E
Sytox Green+ cells
800000
600000
400000
200000
0
0 16 32 40 48
Time (hr)
F
Nsp13-WT
Nsp13-Tet-Mut
Δ1B-Rec1A-Rec2A
Δ1B
Cleaved Casp3/7+ cells
800000
600000
400000
200000
0
0 16 24 32 40
Time (hr)
G
Sytox Green+ cells
100000
80000
60000
40000
20000
0
0 12 24 36
Time (hr)
****
**
*
****
Mock
Nsp13 WT
Nsp13-Tet-Mut
Δ1B
Δ1B-Rec1A-Rec2A
H
+ SARS-CoV-2
Mock
Mock
Nsp13-WT
Nsp13-Tet-Mut
Δ1B-Rec1A-Rec2A
Δ1B
N-protein
GAPDH

**Figure S1**

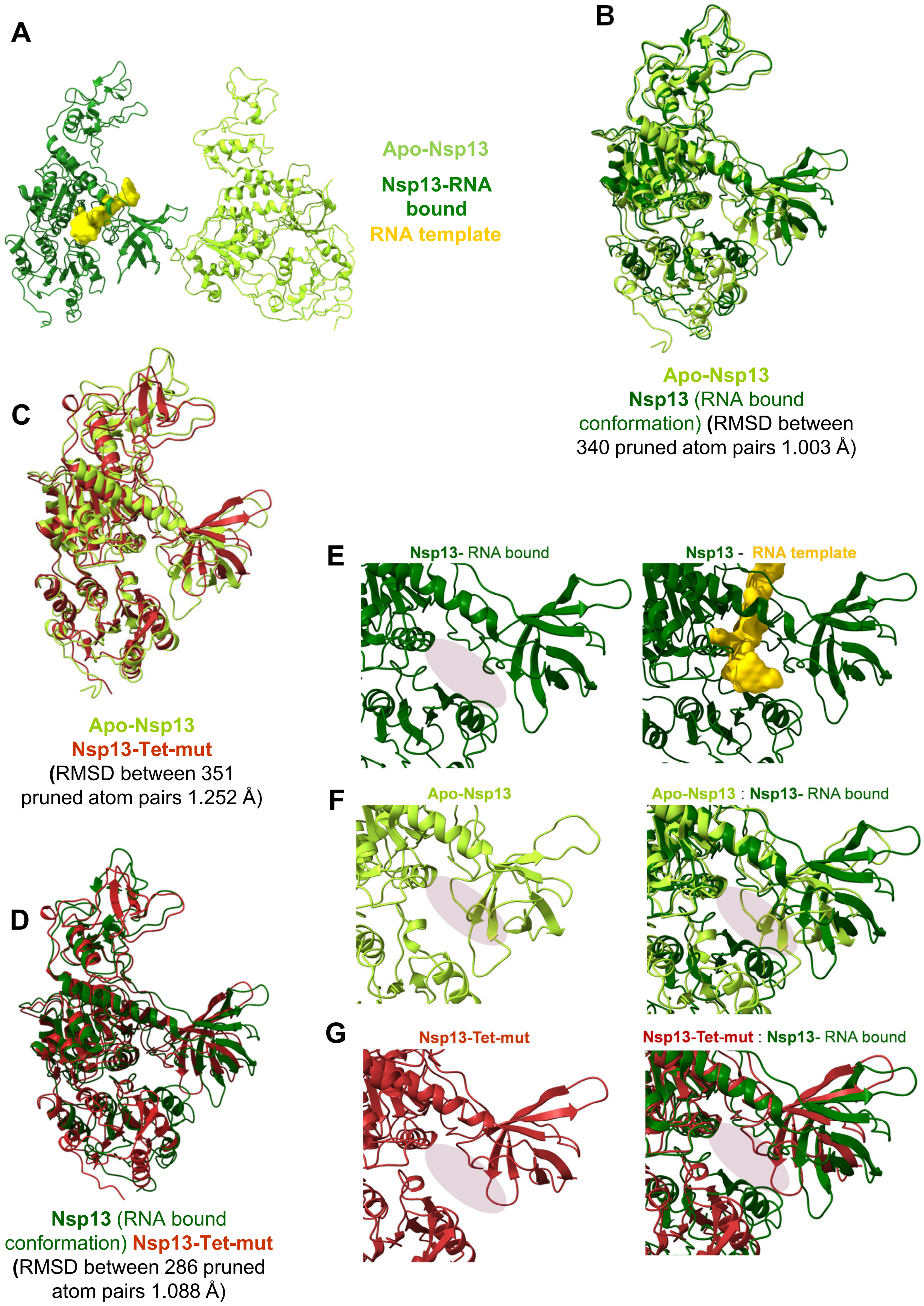
A
Apo-Nsp13
Nsp13-RNA bound
RNA template
B
Apo-Nsp13
Nsp13 (RNA bound conformation) (RMSD between 340 pruned atom pairs 1.003 Å)
C
Apo-Nsp13
Nsp13-Tet-mut
(RMSD between 351 pruned atom pairs 1.252 Å)
D
Nsp13 (RNA bound conformation) Nsp13-Tet-mut
(RMSD between 286 pruned atom pairs 1.088 Å)
E
Nsp13- RNA bound
Nsp13 - RNA template
F
Apo-Nsp13
Apo-Nsp13 : Nsp13- RNA bound
G
Nsp13-Tet-mut
Nsp13-Tet-mut : Nsp13- RNA bound

**Figure S2**

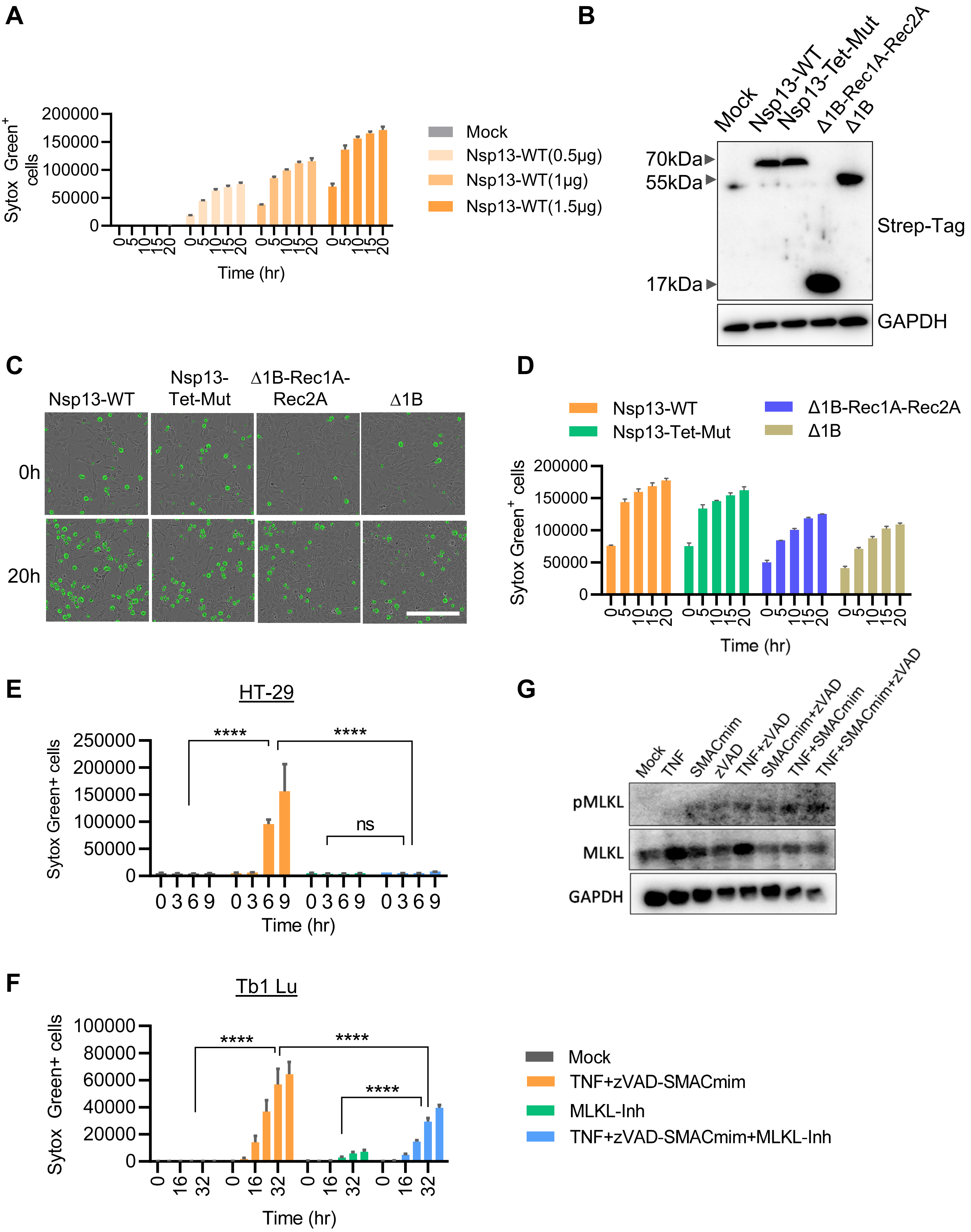
A
Sytox Green+ cells
200000
150000
100000
50000
0
Time (hr)
Mock
Nsp13-WT(0.5μg)
Nsp13-WT(1μg)
Nsp13-WT(1.5μg)
B
Mock
Nsp13-WT
Nsp13-Tet-Mut
Δ1B-Rec1A-Rec2A
Δ1B
70kDa
55kDa
17kDa
Strep-Tag
GAPDH
C
Nsp13-WT
Nsp13-Tet-Mut
Δ1B-Rec1A-Rec2A
Δ1B
0h
20h
D
Nsp13-WT
Nsp13-Tet-Mut
Δ1B-Rec1A-Rec2A
Δ1B
Sytox Green+ cells
Time (hr)
E
HT-29
Sytox Green+ cells
250000
****
ns
Time (hr)
G
Mock
TNF
SMACmim
zVAD
TNF+zVAD
SMACmim+zVAD
TNF+SMACmim
TNF+SMACmim+zVAD
pMLKL
MLKL
GAPDH
F
Tb1 Lu
Sytox Green+ cells
100000
80000
60000
40000
20000
Time (hr)
Mock
TNF+zVAD-SMACmim
MLKL-Inh
TNF+zVAD-SMACmim+MLKL-Inh

**Figure S3**

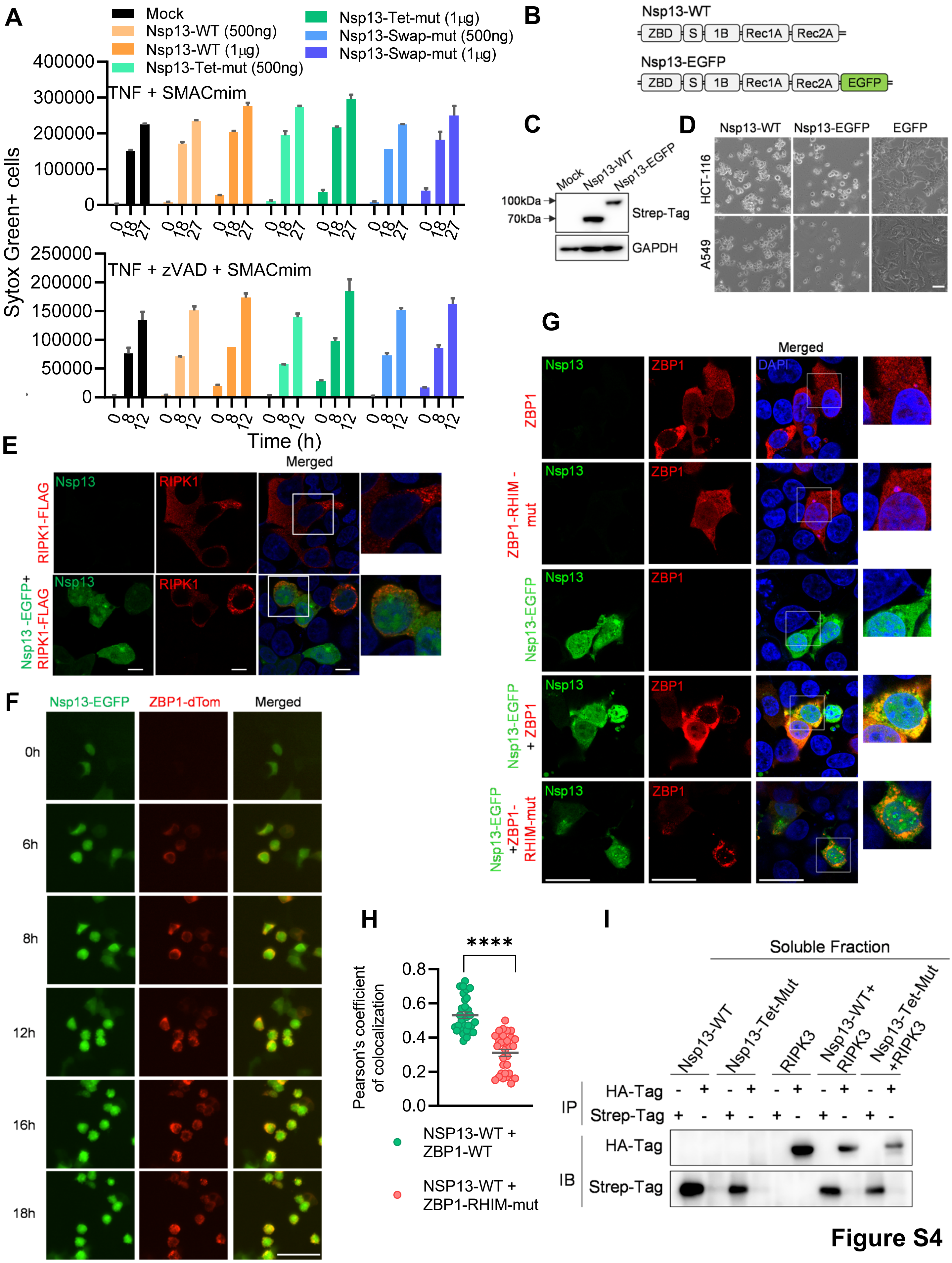
A
Mock
Nsp13-WT (500ng)
Nsp13-WT (1μg)
Nsp13-Tet-mut (500ng)
Nsp13-Tet-mut (1μg)
Nsp13-Swap-mut (500ng)
Nsp13-Swap-mut (1μg)
TNF + SMACmim
TNF + zVAD + SMACmim
Sytox Green+ cells
Time (h)
B
Nsp13-WT
ZBD S 1B Rec1A Rec2A
Nsp13-EGFP
ZBD S 1B Rec1A Rec2A EGFP
C
Mock
Nsp13-WT
Nsp13-EGFP
100kDa
70kDa
Strep-Tag
GAPDH
D
Nsp13-WT
Nsp13-EGFP
EGFP
HCT-116
A549
E
Nsp13
RIPK1
Merged
RIPK1-FLAG
Nsp13 -EGFP+ RIPK1-FLAG
F
Nsp13-EGFP
ZBP1-dTom
Merged
0h
6h
8h
12h
16h
18h
G
Nsp13
ZBP1
DAPI
Merged
ZBP1
ZBP1-RHIM - mut
Nsp13-EGFP
Nsp13-EGFP + ZBP1
Nsp13-EGFP +ZBP1- RHIM-mut
H
Pearson's coefficient of colocalization
****
NSP13-WT + ZBP1-WT
NSP13-WT + ZBP1-RHIM-mut
I
Soluble Fraction
Nsp13-WT
Nsp13-Tet-Mut
RIPK3
Nsp13-WT+ RIPK3
Nsp13-Tet-Mut +RIPK3
IP
HA-Tag
Strep-Tag
IB
HA-Tag
Strep-Tag

**Figure S4**

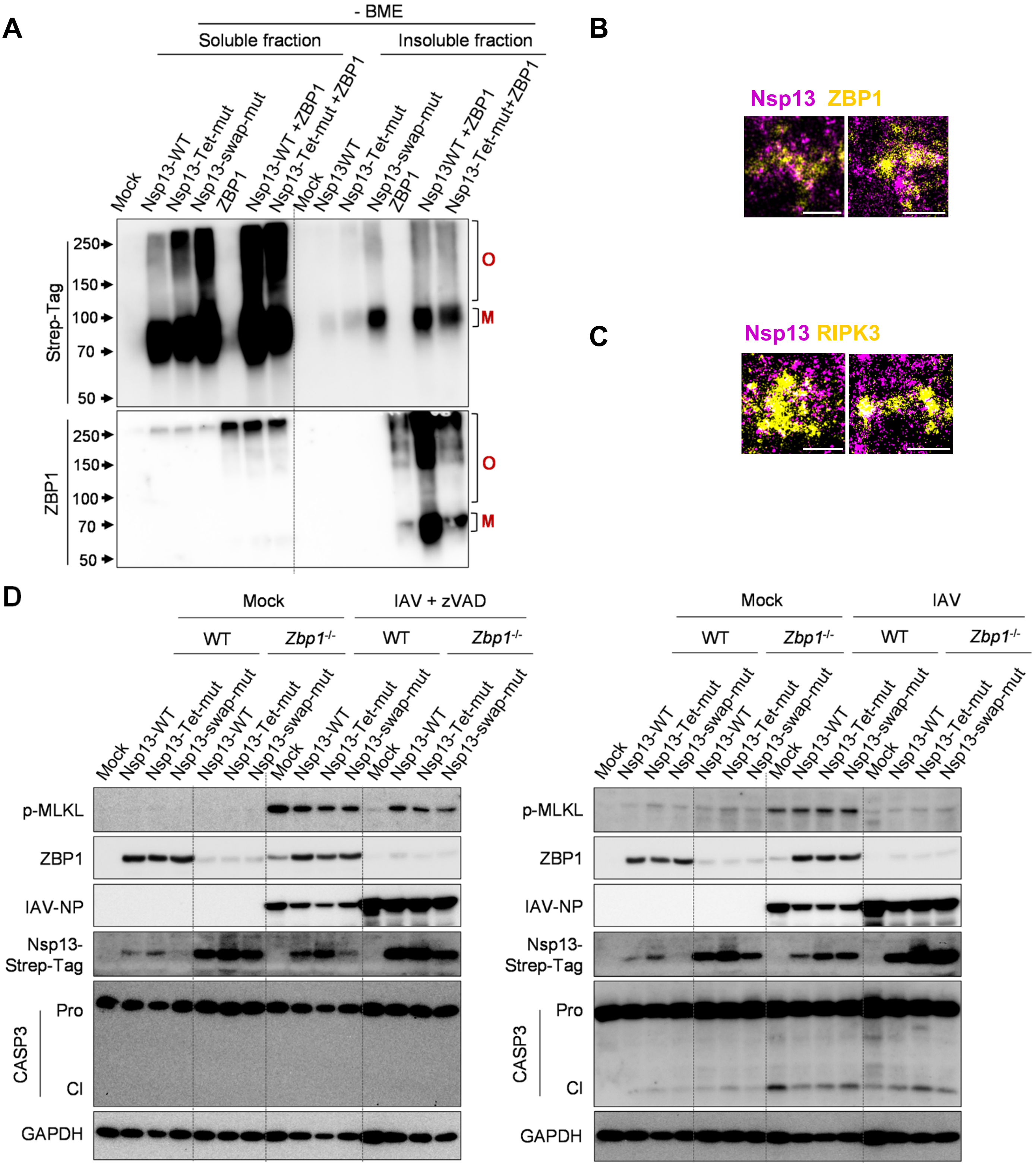
A
- BME
Soluble fraction
Insoluble fraction
Mock
Nsp13-WT
Nsp13-Tet-mut
Nsp13-swap-mut
ZBP1
Nsp13-WT +ZBP1
Nsp13-Tet-mut +ZBP1
Nsp13WT
Nsp13WT +ZBP1
Nsp13-Tet-mut+ZBP1
Strep-Tag
ZBP1
250
150
100
70
50
O
M
B
Nsp13 ZBP1
C
Nsp13 RIPK3
D
Mock
IAV + zVAD
IAV
WT
Zbp1-/-
p-MLKL
IAV-NP
Nsp13-
Strep-Tag
CASP3
Pro
Cl
GAPDH

Figure S5

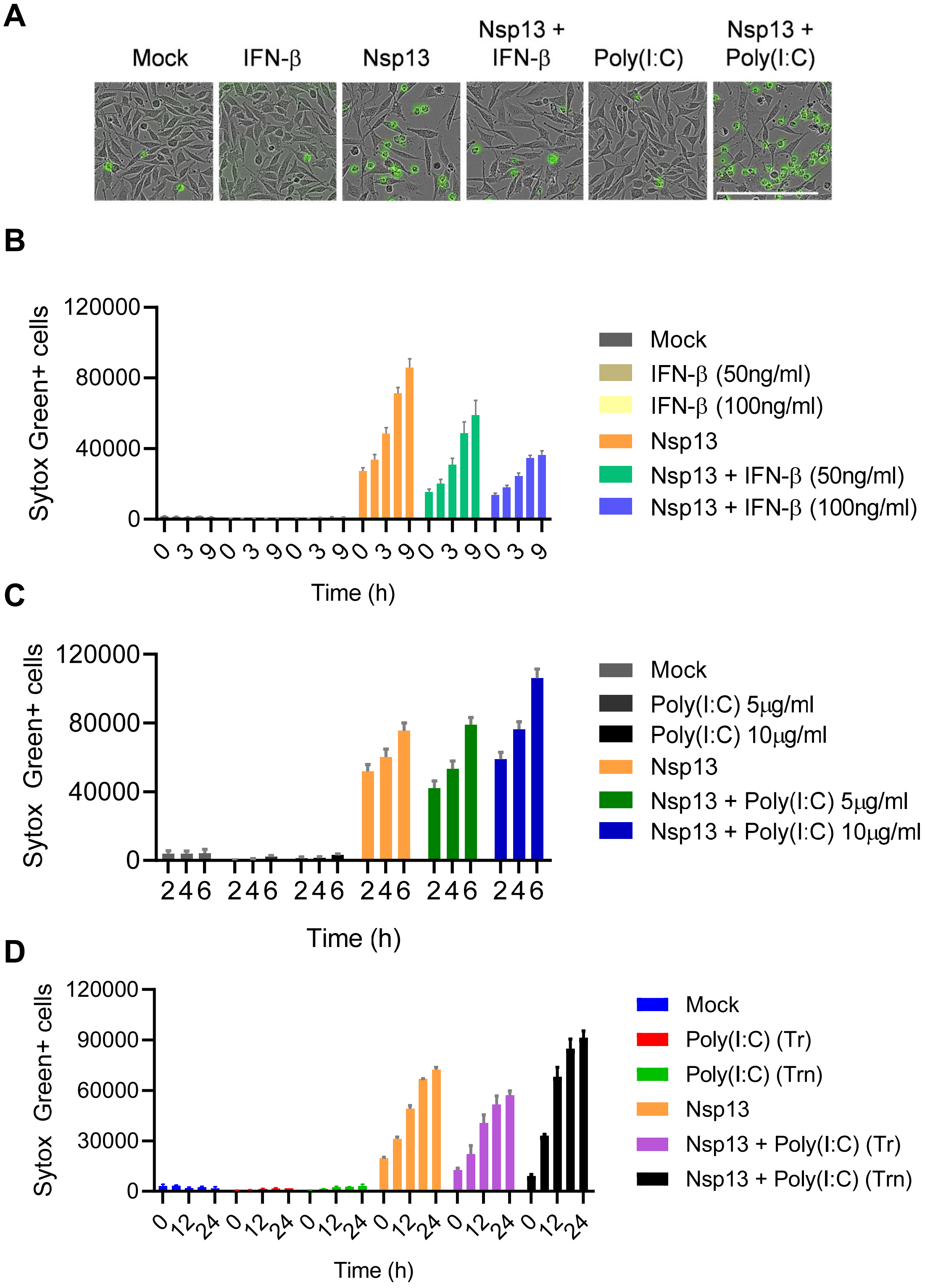
A
Mock
IFN-β
Nsp13
Nsp13 + IFN-β
Poly(I:C)
Nsp13 + Poly(I:C)
B
Sytox Green+ cells
120000
80000
40000
0
0 3 9 0 3 9 0 3 9 0 3 9 0 3 9 0 3 9
Time (h)
Mock
IFN-β (50ng/ml)
IFN-β (100ng/ml)
Nsp13
Nsp13 + IFN-β (50ng/ml)
Nsp13 + IFN-β (100ng/ml)
C
Sytox Green+ cells
120000
80000
40000
0
2 4 6 2 4 6 2 4 6 2 4 6 2 4 6 2 4 6
Time (h)
Mock
Poly(I:C) 5μg/ml
Poly(I:C) 10μg/ml
Nsp13
Nsp13 + Poly(I:C) 5μg/ml
Nsp13 + Poly(I:C) 10μg/ml
D
Sytox Green+ cells
120000
90000
60000
30000
0
0 12 24 0 12 24 0 12 24 0 12 24 0 12 24 0 12 24
Time (h)
Mock
Poly(I:C) (Tr)
Poly(I:C) (Trn)
Nsp13
Nsp13 + Poly(I:C) (Tr)
Nsp13 + Poly(I:C) (Trn)

**Figure S6**

A

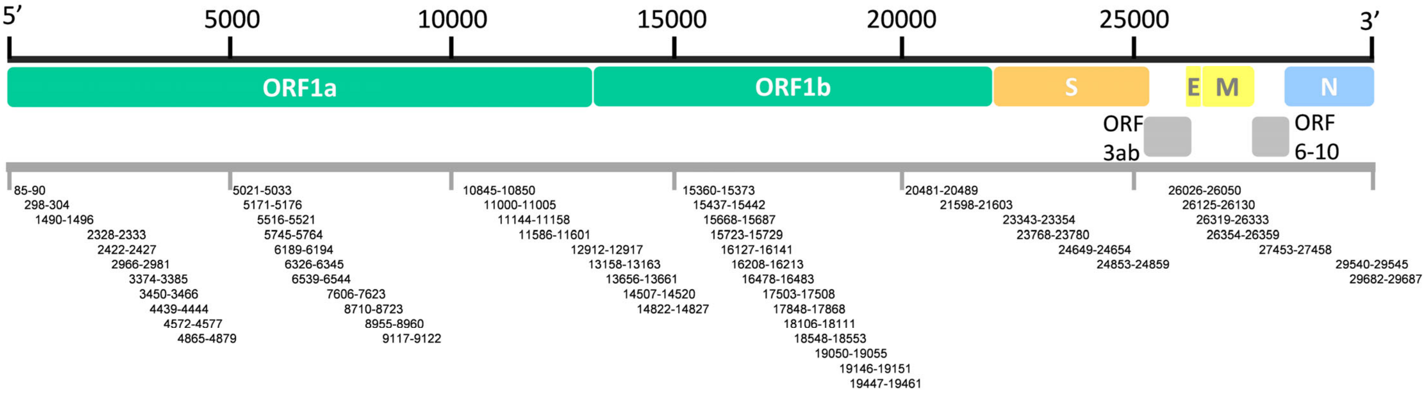


B

| Predicted Z-RNAs | SARS-CoV-2 genome sequence | Sequence location (coding sequence) |
|---|---|---|
| SC2-zRNA-1 | GCACAUCUAGGUUUCGUCCGGGUGUGA | 227-253 (Nsp1) |
| SC2-zRNA-2 | UGAGUGGAGUAUGGCUACAUACUCACUUA | 2959-2987 (Nsp2) |
| SC2-zRNA-3 | CUAGUGAGUACACUGGUAAUUACCAGUGUGGUCACUAU | 5736-5774 (Nsp3) |
| SC2-zRNA-4 | UGAUGGUGGUGUCACUCGUGACAUAGCAUCUA | 8699-8731 (Nsp4) |
| SC2-zRNA-5 | CUGACACACGUUAUGUGCUCAUG | 9114-9136 (Nsp4) |
| SC2-zRNA-6 | UGAUGUUUGUGAAACAUAAGCAUG | 11142-11165 (Nsp6) |
| SC2-zRNA-7 | UGAGUUAACAGGACACAUGUUAGACAUGUAUUCUGUUAUGC UUAC | 16120-16163 (Nsp13) |
| SC2-zRNA-8 | GGCACACUAGAACCAUUUCAAUUCAGUGUGUAGAC | 17479-17516 (Nsp13) |
| SC2-zRNA-9 | UAACAGAUGCGCAAACAGGUUCAUCUAAGUGUGUGUGUUCU GUU | 20459-20504 (Nsp16) |
| SC2-zRNA-10 | GCUGUACUCAACUCAAUUGAGUACAG | 26026-26050 (N protein) |

C

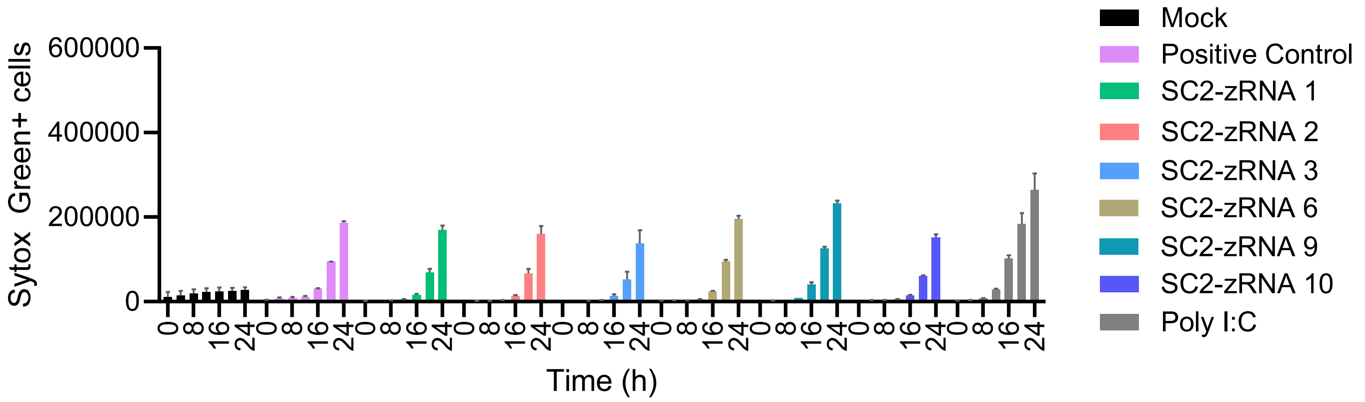


**Figure S7**
